# Neonatal exposures to sevoflurane in rhesus monkeys alter synaptic ultrastructure in later life

**DOI:** 10.1101/2022.03.08.483510

**Authors:** Tristan Fehr, William G.M. Janssen, Janis Park, Mark G. Baxter

## Abstract

Early-life exposure to anesthesia in infant humans and monkeys increases the risk for cognitive and socioemotional impairments. However, the long-term effects of neonatal anesthesia on synaptic ultrastructure have not been thoroughly investigated in primates. We used electron microscopy with unbiased stereological sampling to assess synaptic ultrastructure in the CA1 of the hippocampus and the dorsolateral prefrontal cortex (dlPFC) of female and male rhesus macaques four years after three 4-hour exposures to sevoflurane during the first five postnatal weeks. We counted synapses and measured synapse areas for all synapses and those classified as perforated or nonperforated with spine or dendritic shaft targets. We measured numbers and shapes of mitochondria within presynaptic boutons and calculated vesicle docking rates. In monkeys exposed to anesthesia as infants, synapse areas were reduced in the largest 20% of synapses in CA1 and the largest 5% of synapses in dlPFC, with differential sex effects for the largest 10% of synapses in CA1. Synapse areas were reduced by 7.6% for perforated spinous synapses in CA1, and by 10.4% for nonperforated spinous synapses in dlPFC. Perforated and nonperforated dendritic synapse numbers in CA1 increased by 180% and 63% respectively. Curved mitochondria decreased 25% in CA1 after anesthesia exposure, and dlPFC boutons with 0 mitochondria showed an interaction of anesthesia and sex. These results demonstrate that exposure to anesthesia in infancy can cause long-term structural deficits in primates. These structural changes may be substrates for long-term alterations in the strength and efficiency of synaptic transmission in hippocampus and prefrontal cortex.

**Key points:**

1. Exposure to anesthesia in early life causes lasting cognitive and socioemotional impairments in human and nonhuman primates, but the extent to which early-life exposure to anesthesia alters synaptic ultrastructure in later life has not been known.
2. Four years after rhesus monkeys were given multiple exposures to anesthesia in infancy, the area of spinous synapses was reduced in CA1 and dlPFC, dendritic synapse numbers were elevated in CA1, there were fewer curved presynaptic mitochondria in CA1, and numbers of presynaptic boutons without mitochondria were altered in dlPFC.
3. The long-term ultrastructural changes to synapses and presynaptic mitochondria of rhesus monkeys that were exposed to anesthesia as infants could help explain their behavioral deficits later in life.

## INTRODUCTION

An estimated 500,000 to 1 million children under age 3 in the US are exposed to general anesthesia every year (Ing et al., 2021). Retrospective human epidemiological studies show that early-life exposure to anesthesia can affect subsequent childhood and adolescent behaviors, particularly following multiple exposures. Children exposed to anesthesia early in life are at greater risk for developing learning disabilities (Flick et al., 2011; Ing et al., 2012, 2014; Wilder et al., 2009), socioemotional behavioral problems (DiMaggio et al., 2012), and attention deficit-hyperactivity disorder (Hu et al., 2017; Ing et al., 2020; Sprung et al., 2012; Tsai et al., 2018). Recently, a meta-analysis of prospective cohort studies showed that even a single exposure to anesthetics during early life can confer negative effects on socioemotional behaviors and executive function in children and adolescents (Ing et al., 2021; Jevtovic-Todorovic, 2021).

The long-term behavioral outcomes of childhood exposure to anesthesia have been recapitulated in nonhuman primate models that have ethological and developmental similarity to humans. Rhesus monkeys that were exposed to anesthesia as infants exhibit cognitive deficits (Paule et al., 2011; Talpos et al., 2019) and altered socioemotional behaviors (Coleman et al., 2017; Neudecker, Perez-Zoghbi, & Brambrink, 2021; Neudecker, Perez-Zoghbi, Coleman, et al., 2021) in subsequent years. In the present cohort of rhesus monkeys exposed multiple times to anesthesia as infants, we also found deficits in visual recognition memory at 24-30 months of age (Alvarado et al., 2017) and altered socioemotional behavior characterized by increased reactivity to a social stressor from 6-48 months (Raper et al., 2015, 2018, in prep.).

However, the long-term cellular and molecular mechanisms that accompany behavioral changes in the primates exposed to anesthesia as infants remain insufficiently characterized. In contrast to behavioral outcomes shown over years, neurotoxic effects of neonatal anesthesia in nonhuman primates have been shown on a short-term scale of hours to weeks following anesthesia exposure (Walters & Paule, 2017). A pressing question remains: which neurobiological disturbances that are triggered by exposure to anesthesia in early life persist into the long-term, and could be linked to the long-term behavioral effects seen in primates?

Synapses are one likely substrate for the long-term effects of anesthesia on behavior. Exposure of rodents to anesthesia in infancy cause a deterioration of synaptic signaling, as evinced by suppressed long-term potentiation (LTP) (Guo et al., 2018; Jevtovic-Todorovic et al., 2003; Kato et al., 2013; Schaefer et al., 2020; Sun et al., 2020), inhibited vesicular exocytosis (Baumgart et al., 2015), and altered expression of vesicular docking proteins (Atluri et al., 2019; Xiao et al., 2016). Neonatal anesthesia exposures also alter the overall density of synapses in the hippocampus and subiculum of rodents (Amrock et al., 2015; Ju et al., 2019; Lunardi et al., 2010), and could impact the morphology of remaining synapses. Dendritic spine shapes that are tightly correlated to synapse size and strength (Harris & Weinberg, 2012) are subject to change by early-life anesthesia in the hippocampus and frontal cortex of rodents (Granak et al., 2021). Specifically, early-life anesthesia is linked to the reduction of large mushroom spines (Schaefer et al., 2020) associated with large, stable perforated synapses and LTP (Buchs & Muller, 1996), and the proliferation of smaller, immature spines and filopodia (Briner et al., 2010, 2011; De Roo et al., 2009; Zimering et al., 2016). Together, these findings suggest synaptic changes occur in the long term as well, and are a possible mechanism for the behavioral effects of anesthesia seen years after exposure in infancy.

The ongoing dysfunction of mitochondria is another possible consequence of early-life anesthesia and contributor to behavioral impairments. The health of mitochondria and their contribution to synaptic transmission can be estimated by their shape: straight and curved presynaptic mitochondria have larger active zones and greater vesicle docking rates than donut-shaped mitochondria in monkey dorsolateral prefrontal cortex (dlPFC) (Hara et al., 2014). Likewise, monkeys’ accuracy on a delayed response task is positively correlated with the number of straight mitochondria per bouton and negatively correlated with the number of donut mitochondria per bouton in dlPFC (Hara et al., 2014), linking mitochondrial shape to cognition. After exposure to anesthesia in early life, however, there is a reduction in overall mitochondria density and degradation of the ultrastructure of mitochondria in the hippocampus and subiculum of rodents, concomitant to deficits in cognition and synaptic transmission (Lunardi et al., 2010; Sanchez et al., 2011; Wu et al., 2017). These changes to mitochondria are attributed in part to an elevation in reactive oxidative species (ROS) that impacts their shape, number, and function by disrupting the processes of fission and fusion (Boscolo et al., 2013)—processes that recycle damaged mitochondria and compensate for their energetic dysfunction (Norat et al., 2020). The influx of ROS and the related depolarization of mitochondrial membranes by anesthesia (Bains et al., 2009; Ljubkovic et al., 2007) can also trigger mitochondria morphogenesis from tubular into vase and donut shapes, both through failures of fusion (X. Liu & Hajnóczky, 2011), and independent of fusion processes (Miyazono et al., 2018). In turn, donut and other degraded mitochondria contribute to the ongoing generation of ROS (Ahmad et al., 2013). Protecting mitochondria from ROS not only prevents anesthesia’s deleterious effects on mitochondria density and ultrastructure, but also prevents cognitive impairments caused by anesthesia exposure (Boscolo et al., 2012; Wu et al., 2017). Abnormal mitochondrial numbers and shapes within presynaptic boutons of monkeys years after anesthesia exposure in infancy could therefore indicate a chronic dysfunction of mitochondria related to deficits in synaptic transmission and behavior.

We chose the CA1 of the hippocampus as one region in which to investigate the long-term changes to synapse ultrastructure caused by neonatal exposure to anesthesia. Hippocampal damage is behaviorally suggested in the present cohort because impairments in visual recognition and disrupted socioemotional behaviors follow neonatal hippocampal lesions (Raper et al., 2017; Zeamer et al., 2010). Further, the effects of early-life anesthesia in the primate CA1 may mirror synaptic impacts of neonatal anesthesia in the rodent CA1, including short- and long-term downregulation of LTP, impaired vesicle docking, and synapse loss (Atluri et al., 2019; De Roo et al., 2009; Schaefer et al., 2020; Wu et al., 2016; H. Zhang et al., 2015).

We also chose to probe layer III of the dorsolateral prefrontal cortex (dlPFC). Neurons in the dlPFC are critical to emotional regulation and working memory (Smits et al., 2020), and synaptogenesis in the dlPFC layer III in the first 1.5 years of life (Anderson et al., 1995) coincides with the timeframe for anesthesia exposure in our cohort, opening a potential window of vulnerability to anesthetic insults. Likewise, the development and function of dlPFC neurons are dependent on hippocampal integrity (Bertolino et al., 1997; Saunders et al., 1998), which could be compromised by anesthesia.

We hypothesized that repeated exposure of monkeys to the common pediatric anesthetic sevoflurane, in three 4-hour sessions over the first five postnatal weeks, would result in altered synapses and mitochondria within CA1 of the hippocampus and the dlPFC four years after exposure. We measured the synapse areas, synapse density, and rates of vesicle docking across all synapses in these regions, and then measured the areas and numbers of synaptic subtypes characterized by their targets and complexity. Additionally, we measured the total density, shape, and frequency of mitochondria in presynaptic boutons. We report that repeated anesthesia exposure during infancy in nonhuman primates differentially impacted the synaptic ultrastructure in area CA1 of the hippocampus and in the dlPFC during adolescence, with sex interactions in CA1 and specific effects for synapse subtypes. We also report anesthesia exposure in early life changed the prevalence of mitochondrial shapes in CA1 and interacted with sex to affect the number of mitochondria per presynaptic bouton in dlPFC.

## MATERIALS & METHODS

### Monkey anesthesia and behavior

As previously described (Raper et al., 2015), infant rhesus macaques in the anesthesia group (n = 5 female, n = 5 male) received a 4-hour exposure to ∼2.5% sevoflurane on ∼P7 (range P6-10), two weeks later at ∼P21, and two weeks later again at ∼P35 for a total of 3 anesthetic exposures between ∼P7-35. Monkeys in the control group (n = 5 female, n = 5 male) received a brief maternal separation for ∼30 min following the same time schedule as the anesthesia group.

After removal from their dam, all infants received a brief neurological exam. At this point, anesthesia group monkeys were mask-induced with sevoflurane (from 2 vol % to effect, maximum 8 vol % in 100% O_2_), intubated, and catheterized for IV fluids. Sevoflurane was administered for 4 hours, with monitoring of vital signs, depth of anesthesia and blood gasses. Results of physiological monitoring during anesthesia were consistent with normal physiology, with no indications of hypoxemia, hypercapnia, or hypotension. Upon complete recovery usually within 20-30 min, the infant was returned to its dam. For control group subjects, the maternal separation procedure consisted of the neurological exam and a period of handling with a duration of separation that matched that of the period of conscious separation experienced by the experimental group. On average, control infants experienced 30-40 min of maternal separation, and were returned to their dam. Mother-infant interactions after these separations did not differ between groups, indicating that the separations involved in anesthesia exposure did not alter mother-infant bonding that might have impacted later cognitive or socioemotional behavior (Raper et al., 2016).

At 6, 12, 24, and 48 months, monkeys were tested in a human intruder paradigm (Raper et al., 2015, 2018, in prep.) and were also tested in a visual paired comparison test of recognition memory at 6, 12, and 24 months (Alvarado et al., 2017). Home environment observations were also collected (unpublished).

### Brain collection and preparation for electron microscopy

At ∼48 months of age, monkeys were perfused following previously described methods (Hara et al., 2014). Monkeys were deeply anesthetized with ketamine hydrochloride (25 mg/kg) and pentobarbital (20–35 mg/kg, i.v.), intubated, and mechanically ventilated to prevent hypoxia and ischemia. Monkeys were transcardially perfused with cold 1% paraformaldehyde and 0.125% glutaraldehyde in 0.1 M phosphate buffer (PFA/PB; pH 7.2) for 2 min, followed by 4% (wt/vol) PFA/PB at 250ml/min for 10 min and 100ml/min for 50 min.

After perfusion, tissue was postfixed for 6 hours in 4% PFA/PB with 0.125% glutaraldehyde, washed in PB. Coronal blocks containing the dorsolateral prefrontal cortex or hippocampus were sectioned at 50- and 400-μm thicknesses on a vibratome (Leica) for light and electron microscopy, respectively. Specifically, 12 50-μm sections and one 400-μm section were taken in every mm of brain tissue, with a random placement of the first 400-μm section within the series of 50-μm sections to allow for stereologic sampling. From the 400-μm sections, we generated two sets of tissue blocks for EM analysis.

Tissue sections were blocked to identify hippocampal CA1 at a mid-anterior/posterior level from the body of the hippocampus posterior to the uncus, as well as the dlPFC spanning the principal sulcus (Brodmann’s area 46). Tissue was washed in sodium cacodylate buffer, placed in 1% osmium tetroxide/double distilled water (ddH2O) for 1 hour, washed with ddH2O, and stained en bloc in cold 2% uranyl acetate/ddH2O for 1 hour in the dark. After a ddH2O wash, tissue was dehydrated through an ascending ethanol/ddH2O series, washed 10 min in 1:1 ethanol:propylene oxide (PO), 10 min in PO, and infiltrated with EPON resin (Electron Microscopy Sciences, EMbed 812 Kit): ascending resin:propylene oxide with pure resin overnight and again the next morning. Blocks were embedded in BEEM capsules and placed in a vacuum for 72 hours at 60°C.

In preparation for ultrastructural analysis, tissue from stratum radiatum of CA1 and layer III (250-350 µm from the layer I-II intersection) of dlFPC was semi-thin sectioned at 1 µm on a Leica UC-7 ultramicrotome, and stained with toluidine blue to further identify regions of interest for ultrathin (70 nm) serial sections. Ribbons of 15-20 ultrathin (70 nm) serial sections were cut using a Diatome diamond knife (Electron Microscopy Sciences), mounted on Formvar-coated slot grids (Electron Microscopy Sciences), and counter-stained with uranyl acetate and lead citrate.

### Imaging on electron microscope

All electron microscopy imaging, image processing, and image analysis was conducted by an experimenter blind to experimental condition and sex of the monkeys. Monkeys were assigned a random alpha-numeric code for tracking purposes during imaging and image analysis that contained no information about their sex or anesthesia group. This code was broken only after all image analyses had been completed. One control female monkey and one anesthesia female monkey were excluded from further analysis due to an unresolvable possible confusion of their tissue blocks during initial processing. Thus, samples from 9 control monkeys (n = 5 male, n = 4 female) and 9 anesthesia monkeys (n = 5 male, n = 4 female) continued forward into further analysis.

Slot grids containing ribbons of at least 15 serial sections from the region of interest were imaged on a Hitachi H-7000 TEM microscope (Hitachi High Technologies America, Inc.) with an AMT Advantage CCD camera (Advanced Microscopy Techniques) at 12,000x. Briefly, 5 non-overlapping image series of 15 sequential sections were captured per animal. Care was taken to avoid blood vessels, cell bodies, and post-staining artifacts in the image fields. Imaging was conducted in tandem with post-exposure image processing in Adobe Photoshop (CS5) to maximize field alignments, sharpness, and brightness between sections. Image field area was 83.09 µm^2^, and section thickness was 70 nm per section, for a section volume of 5.82 µm^3^. The total volume per 15-section series was 87.25 µm^3^.

### Image processing and quantitative analyses

Each image series of 15 sequential sections from the CA1 stratum radiatum or dlPFC layer III was imported into the open-source software Reconstruct (https://synapseweb.clm.utexas.edu/software-0) to create reconstructions of brain tissue for feature analysis in three dimensions. Section images were sequentially aligned to each other using the manual linear alignment tool beginning at the middle, or 8^th^, section and moving to either end of the 15-section series. CA1 and dlPFC measurements were tabulated separately.

#### Synapse areas

Synapses were identified by presence of an electron-rich postsynaptic density. Each 15-section series was subdivided into 3 non-overlapping series of 5 sections. For each of these 5-section subseries, the lengths of synapses identified on the middle, or 3^rd^, section were traced to completion across the sections using Reconstruct’s z-tracing tool. Synapse areas were calculated by multiplying synapse lengths by the total thickness of the sections they crossed (.070 µm * number of sections). Synapses that intersected the x- and y-limits of the sections were excluded from analysis. Areas of n = ∼250-330 synapses in dlPFC and n = ∼350-450 synapses in CA1 were calculated per animal.

#### Synapse area classifications by type

Synapses were also classified by their postsynaptic target: dendritic spines (spinous synapses) or dendritic shaft (dendritic synapses); and synapse complexity: perforated morphology or nonperforated (macular or continuous) morphology. Perforated synapses were identified by a discontinuity in the postsynaptic density. Once synapses were classified according to their targets and complexity, synapse areas and total synapse number in each category per 5-section series (29.08 µm^3^) were calculated. Examples of perforated spinous (**Figure 1A**), perforated dendritic (**Figure 1B**), nonperforated spinous (**Figure 1C**), and nonperforated dendritic (**Figure 1D**) synapses are shown.

**Figure 1.**
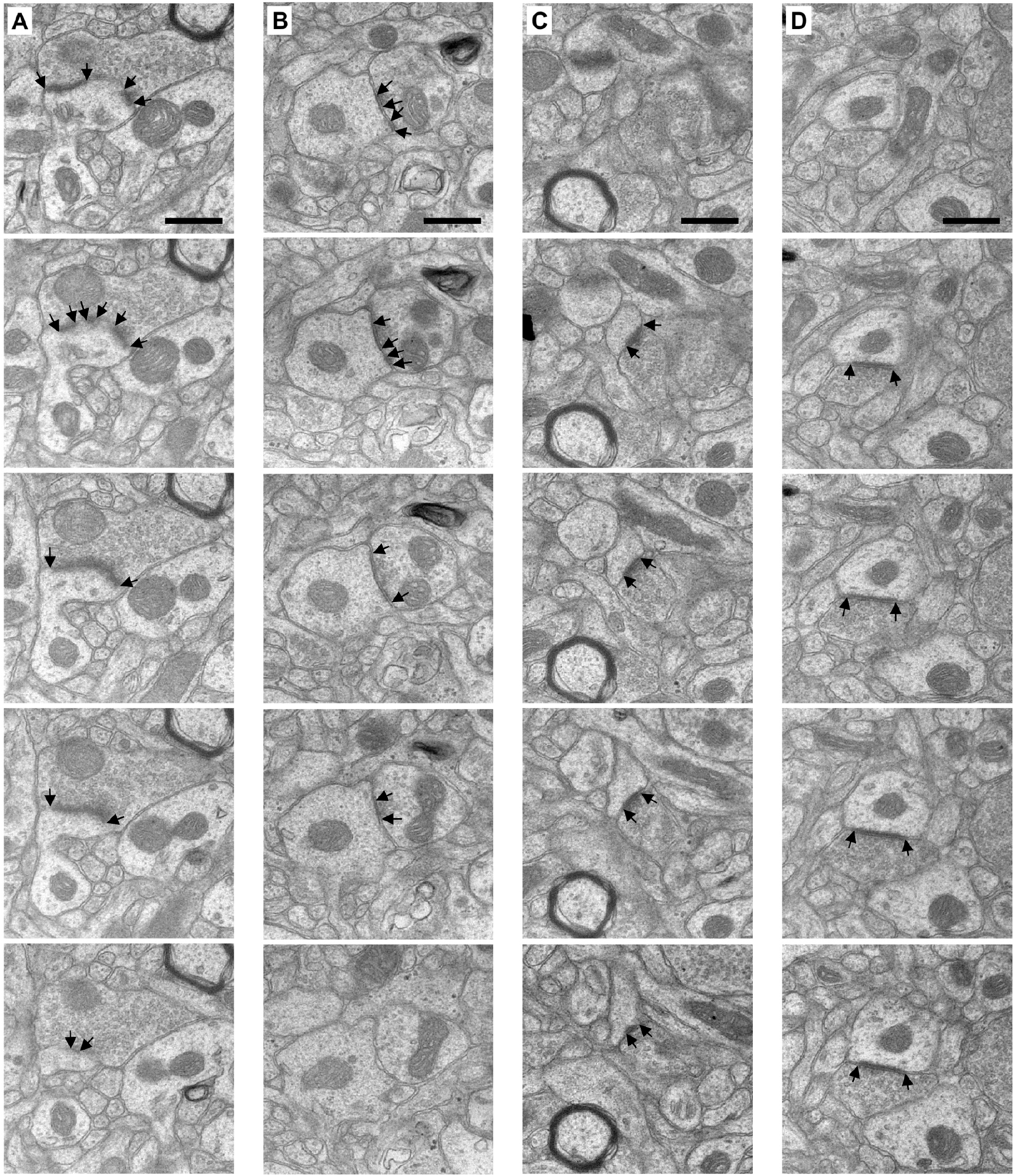
From left to right, serial sections of a perforated spinous (*A*), perforated dendritic (*B*), nonperforated spinous (*C*), and nonperforated dendritic synapse (*D*). Black arrows denote the boundaries of the synapse. Scale bar is 500 nm. Serial sections are 70 nm thick.

#### Synapse density with optical disector

Synapse density was measured using an optical disector approach. 5 separate pairs of adjacent section images were selected from each of three 15-section series, for a total of 15 pairs per animal. For each pair of sections, only the synapses that were unique to each section and did not appear on the adjacent section were counted. The number of unique synapses at the leading edge in each section was divided by the section volume of 5.82 µm^3^ to calculate synapse density for n=30 sections per animal.

#### Vesicle docking

We used measures of vesicle docking to estimate synaptic activity in our regions of interest. We used the program SynBin (Adams et al., 2001) to define the boundaries of presynaptic boutons and the lengths of synapses, and to mark vesicles therein for quantification. Vesicles in axonal boutons are each ∼35 nm and docked vesicles are often smaller than docked vesicles (Harris & Weinberg, 2012). Therefore, we estimated that actively docking vesicles would appear within 30 nm of the presynaptic membrane, while vesicles that are 30-60 nm away from the membrane represent vesicles that are not actively docking. For each of 10 synapses per section, we counted the number of vesicles 0-30 nm and 30-60 nm from the inner edge of the presynaptic membrane, defined respectively as the docking and pre-docking zones. We calculated vesicle docking as the ratio of the number of docked vesicles to the sum of docked and pre-docked vesicles for each synapse. In each of the 5 series of 15 sections per animal, we performed analyses on 3 sections that were separated by at least 350 nm (5 sections) to avoid sampling the same synapses, for a total n=150 synapses analyzed per animal per region and n=2,700 total synapses analyzed per region.

#### Presynaptic mitochondria density, number per bouton, and morphology

We assessed the morphology and number of mitochondria within presynaptic boutons across all 5 of the 15-section series per animal, as previously described (Hara et al., 2014). Presynaptic boutons were the focus of mitochondrial analyses at the synapse. To start, we identified presynaptic boutons transected by the middle, or 8^th^, section of each series by the presence of 3 or more synaptic vesicles within their enclosed membranes. We followed each bouton to completion across sections, up and down the z-axis of the 15-section series. In order to not preferentially exclude large boutons, we included boutons that intersected the top (1st) and bottom (15th) sections of the series in the analysis [large boutons comprise an estimated 6-14% of all boutons analyzed (Hara et al., 2016)]. Boutons that were intersected by the x and y axes were excluded from analyses. All measurements of mitochondria pertain only to those within the presynaptic boutons identified in this manner.

Overall mitochondria density was calculated per 15-section series as the sum of mitochondria in presynaptic boutons per 87.25µm^3^ series volume. Likewise, the number of mitochondria within each bouton was counted, and categorized as 0, 1, 2, or 3 or more mitochondria. Finally, mitochondria within each presynaptic bouton were classified and counted within each of three morphological categories: straight, curved, or donut-shaped (toroidal) (**Figure 2A-C**). Curved mitochondria had at least one ≤90° bend, donut mitochondria formed ring shapes, and all other mitochondria were classified as straight.

**Figure 2.**
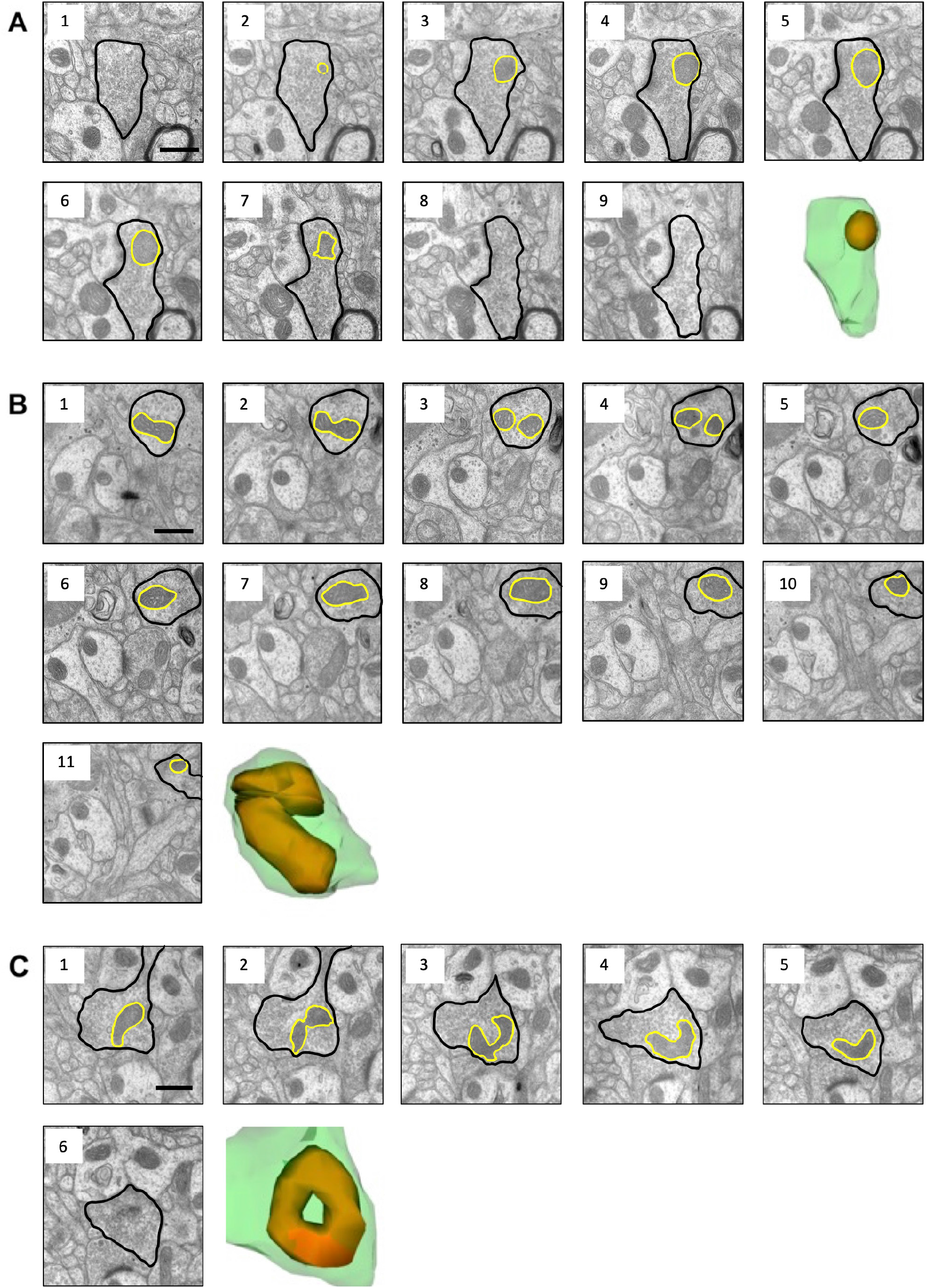
Serial electron microscope images and 3D reconstructions of a straight (*A*), curved (*B*), and donut (*C*) mitochondrion. In electron microscope images, presynaptic bouton is outlined in black and mitochondrion is outlined in yellow. In reconstructions, presynaptic bouton is green and mitochondrion is orange. Scale bars are 500 nm. Serial sections are 70 nm thick.

### Statistical analyses

All statistical analyses were performed separately for CA1 and for dlPFC using the open-source software R (R Core Team, 2020). All descriptive data are reported as the mean ± standard deviation. A complete table of statistical results is included in Supplemental Table 1.

#### Synapse areas, synapse density, vesicle docking, overall mitochondria density

Analyses of synapse areas, measurements of synapse density, vesicle docking ratios, and the overall density of mitochondria in presynaptic boutons were calculated by applying linear mixed models with random effects from the lme4 package (Bates et al., 2015) to case-level data, with fixed effects of group, sex, and their interaction, and random effects of case. Additional analysis of synapse area means by quantile was carried out by calculating estimated marginal means of group by quantile and group by sex by quantile. Homogeneity of variance for synapse density data was evaluated using Bartlett’s test.

#### Counts of mitochondria shapes and numbers per bouton

The counts of: mitochondria classified by shape, the number of mitochondria per bouton across the four numerical categories, and the number of mitochondria per bouton per each count category were analyzed with a hierarchical series of general linear mixed models with random effects using the glmer function of the lme4 package. Poisson distributions were used because the data were counts. Fit of the Poisson distributions to the data was confirmed using the simulateResiduals and testOutliers functions of the DHARMa package (Hartig, 2022). Group, sex, and group by sex interaction contributions to the observed variance were evaluated using hierarchical Wald chi squared tests.

#### Counts of synapse types

Both a full model of synapse counts across synapse types, classified by postsynaptic target and synapse complexity, and models of synapse counts per each synapse type were analyzed with a hierarchical series of general linear mixed models with random effects using the glmer function of the lme4 package. Poisson distributions were used because the data were counts; fit of the Poisson distributions to the data was confirmed using the simulateResiduals and testOutliers functions of the DHARMa package in R. Group, sex, synapse type, and their interactions’ contributions to the observed variance were evaluated using hierarchical Wald chi squared tests.

#### Areas of synapses by type

A full model of synapse areas sorted by postsynaptic target and synapse complexity was analyzed with a linear mixed model with random effects. Wald chi square tests were used to compare a series of hierarchical linear mixed models with random effects to determine group, sex, and group by sex interaction effect contributions to statistical variance across synapse types. Analyses of synapse areas classified by type were carried out using linear mixed models with random effects.

## RESULTS

### Early-life anesthesia exposures affect the areas of the largest CA1 and dlPFC synapses four years later

First, we evaluated the long-term effects of anesthesia on synapse size by measuring synapse areas. In the CA1, monkeys that were exposed to anesthesia as infants showed an overall 8.9% reduction in mean synapse area compared to control monkeys (control 0.32 ± 0.02, anesthesia 0.29 ± 0.02 µm^2^, F(1, 13.98) = 10.68, p = 0.006) (**Figure 3A**). To explore the differences in the distributions of the hundreds of synapse areas we evaluated, we sorted synapse areas in ascending order by size and divided the resulting lineup into 20 bins (quantiles) that each comprised 5% of the total number of synapse area measurements. We found a significant interaction of treatment, sex, and quantile (F(19, 266) = 2.77, p = 0.0002). Pairwise comparisons of control and anesthesia groups showed synapses of monkeys in the anesthesia group were significantly reduced in the 17^th^ (p = 0.02), 18^th^ (p < 0.004), 19^th^ (p = 0.003), and 20^th^ quantiles (p < 0.001) (**Figure 3B**).

**Figure 3.**
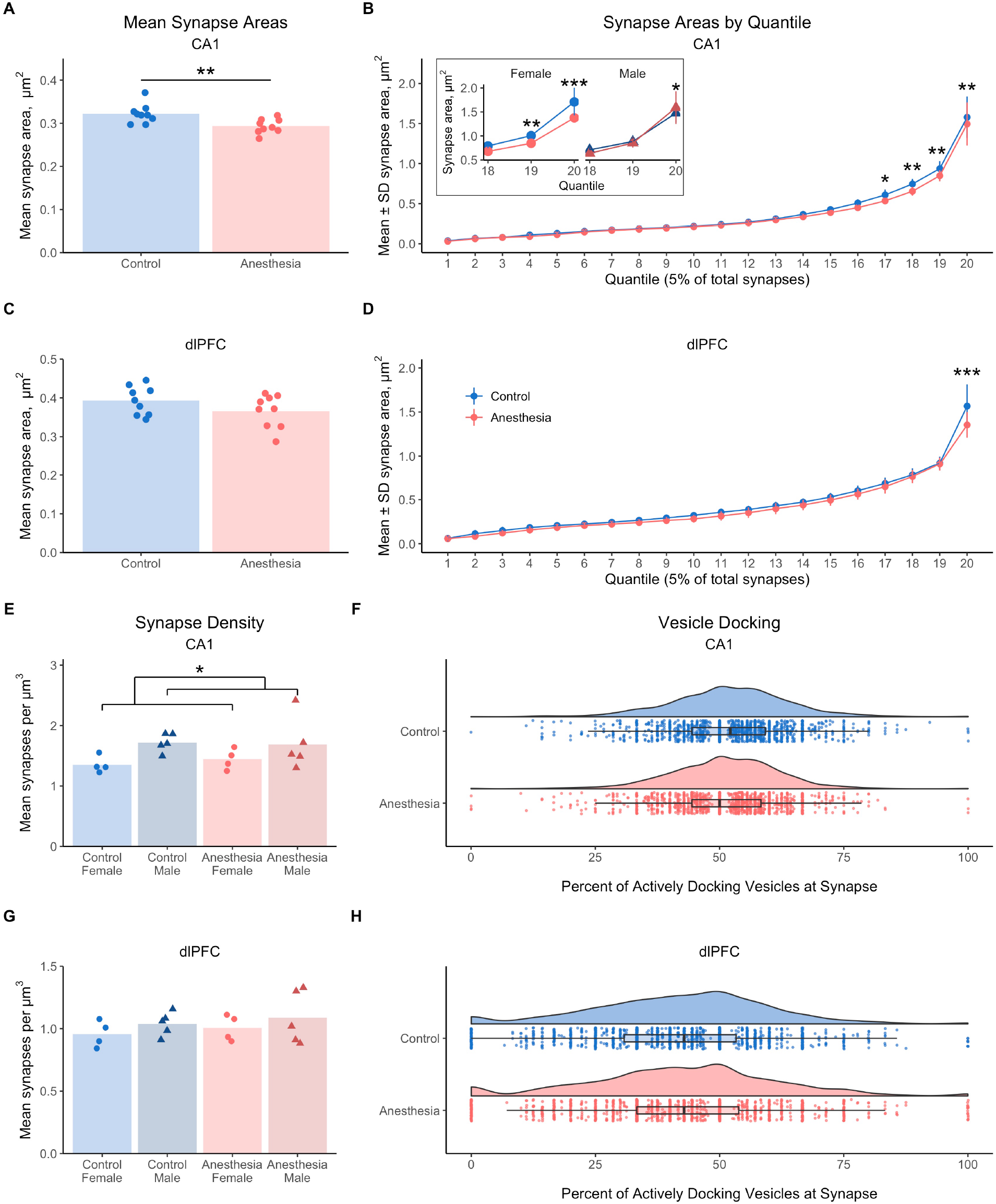
Monkeys exposed to anesthesia as infants show lasting changes to the areas of the largest synapses, but not synapse density or vesicle docking, in CA1 and dlPFC. (*A*) Mean synapse areas are reduced 8.9% in CA1 of monkeys exposed to anesthesia as infants (F(1, 13.98) = 10.68, p = 0.006). (*B*) In CA1, the largest 20% of synapse areas are smaller in monkeys exposed to anesthesia, shown in the 17^th^ (p = 0.02), 18^th^ (p < 0.004), 19^th^ (p = 0.003), and 20^th^ quantiles (p < 0.001). (*B inset*) Females had lower synapse areas associated with anesthesia exposure in the 19^th^ (p < 0.005) and 20^th^ quantiles (p < 0.0001), and males had higher synapse areas in the anesthesia group in the 20^th^ quantile (p = 0.036). (*C*) Mean synapse areas are not affected by anesthesia in dlPFC (F(1, 14.06) = 1.96, p = 0.18). (*D*) The largest 5% of synapses (20^th^ quantile) are reduced in the dlPFC by 13.6% in monkeys exposed to anesthesia as infants (p < 0.0001). (*E*) Synapse density in CA1 was 22% greater in males than females (F(1, 14) = 5.95, p = 0.029), but is not affected by anesthesia (F(1, 14) = 0.06, p = 0.81) or an anesthesia by sex interaction (F(1, 14) =0.23, p = 0.64). (*F*) The proportion of docking vesicles in CA1 synapses was not significantly impacted by early-life exposure to anesthesia (F(1, 14) = 0.88, p = 0.37). (*G*) Synapse density in dlPFC was not affected by anesthesia (F(1, 14) = 0.53, p = 0.48), sex (F(1, 14) = 1.50, p = 0.24), or their interaction (F(1, 14) = <0.0001, p = 0.997). (*H*) The proportion of docking vesicles in dlPFC synapses was not affected by anesthesia (F(1, 13.98) = 0.45, p = 0.51). (*A-H*) Unexposed controls shown in blue, monkeys exposed to anesthesia shown in red. (*A, C, E,* & *G*) Individual points indicate individual monkey means. (*B inset, E,* & *G*) Females are lighter shades with circles, males are darker shades with triangles. (*F* & *H*) Kernel distribution graphs show the relative spread of the frequency of areas, box & whisker plots show the range and quartiles of areas, and individual points are individual synapses. * = p < 0.05, ** = p < 0.01, *** = p < 0.001.

When we included sex as a factor in our CA1 pairwise comparisons (**Figure 3B, inset**), we found that synapse areas in the 19^th^ quantile were lower in females exposed to anesthesia compared to female controls (1.01 ± 0.06 controls, 0.85 ± 0.03 µm^2^ anesthesia, p < 0.005), but not significantly different between males exposed to anesthesia and male controls (0.88 ± 0.08 controls, 0.85 ± 0.09 µm^2^ anesthesia, p = 0.87). Synapse areas in the 20^th^ quantile were lower in females exposed to anesthesia compared to control females (1.71 ± 0.30 control, 1.38 ± 0.05 µm^2^ anesthesia, p < 0.0001), but greater in males exposed to anesthesia compared to control males (1.48 ± 0.20 control, 1.59 ± 0.34 µm^2^ anesthesia, p = 0.036). No other treatment, sex, or quantile effects were observed (p-values > 0.05). These results indicated that while only the synapses with the largest areas in CA1 showed long-term effects of neonatal anesthesia, the direction and magnitude of these effects differed between females and males.

In the dlPFC, we found no significant effects of treatment or sex on mean synapse areas (p-values > 0.15) (**Figure 3C**). However, when synapse areas were sorted into 20 equally sized quantile bins, we observed a significant treatment by quantile interaction (F(19, 266) = 2.81, p = 0.0001). Pairwise comparisons of control and anesthesia groups across quantiles revealed that synapse areas in the 20^th^ quantile were 13.6% lower in monkeys exposed to anesthesia as infants compared to control monkeys (1.57 ± 0.25 control, 1.35 ± 0.15 µm^2^ anesthesia, p < 0.0001) (**Figure 3D**). All other treatment, sex, and quantile effects were not significant (p values > 0.05). Therefore, only the largest synapses in the dlPFC showed long-term impacts of anesthesia exposures in infancy.

### Exposures to anesthesia in early life do not affect synapse density or vesicle docking

We did not find significant effects of anesthesia treatment or treatment by sex interactions on CA1 synapse density (p-values > 0.60), but males showed a 22% greater synaptic density compared to females (females 1.40 ± 0.15, males 1.71 ± 0.31 synapses/µm^3^, F(1, 14) = 5.95, p = 0.029) (**Figure 3E**). Synapse density in dlPFC did not show any effects of treatment or sex (p-values > 0.20) (**Figure 3G**). In sum, synapse density in CA1 and DLPFC was not altered in the long term by neonatal anesthesia exposures.

We measured the proportion of docking vesicles in anesthesia and control monkeys to estimate the long-term impacts of neonatal anesthesia on the baseline synaptic activity in CA1 and dlPFC. However, the proportions of docking vesicles were not significantly affected by anesthesia treatment, sex, or their interaction in CA1 and dlPFC (p-values > 0.15) (**Figures 3F&H**). In sum, we did not find evidence that early-life anesthesia had a long-term impact on baseline vesicle docking in CA1 or dlPFC, suggesting that baseline presynaptic activity is unlikely to be affected by anesthesia exposures years later.

### Impacts of early-life anesthesia on synapses classified by target and complexity

After finding evidence of long-term alterations to the largest synapses in CA1 and dlPFC by exposures to anesthesia during infancy, we next asked whether these changes were specific to synapse type. One element we used to classify synapse types was postsynaptic target, because the size of axodendritic synapses tends to be larger than axospinous synapses in the CA1 of humans (Montero-Crespo et al., 2020). Another synaptic element we used to classify synapse types was synapse complexity: perforated synapses with complex shapes are larger on average than nonperforated synapses with simple disc shapes (Ganeshina et al., 2004b). We hypothesized that the long-term effects of anesthesia on large synapses would translate to differential impacts of anesthesia on synapse numbers and areas, dependent on whether the synapses target dendritic spines or dendritic shafts—which we refer to respectively as “spinous” and “dendritic” synapses—and whether the synapses have perforated or nonperforated morphology.

### Early-life anesthesia boosts long-term numbers of dendritic synapses in CA1

When we counted synapses in the CA1 classified by targets and complexity, we found significant interactions of anesthesia treatment by synapse type (*χ*^2^(3) = 33.89, p = 2.10e^-7^) and sex by synapse type (*χ*^2^(3) = 10.40, p = 0.015). We also found a trend toward an interaction of treatment by sex by synapse type (*χ*^2^(3) = 7.60, p = 0.056). Further analyses showed that the numbers of perforated and nonperforated dendritic synapses were significantly altered by anesthesia exposures. The mean number of perforated dendritic synapses was 180% greater in monkeys exposed to anesthesia as infants compared to controls (controls 0.11 ± 0.32 synapses, anesthesia 0.31 ± 0.60 synapses, *χ*^2^(1) = 6.11, p = 0.013) (**Figure 4A**). The mean number of nonperforated dendritic synapses was 63% greater in anesthesia-exposed monkeys than unexposed controls (controls 0.96 ± 1.13 synapses, anesthesia 1.56 ± 1.43 synapses, *χ*^2^(1) = 6.02, p = 0.014) (**Figure 4C**). There were no additional treatment, sex, or interaction effects for dendritic or spinous synapses in CA1 (p-values > 0.20). These results demonstrated that early-life exposures to anesthesia caused a long-term, selective increase in numbers of dendritic synapses in the CA1.

**Figure 4.**
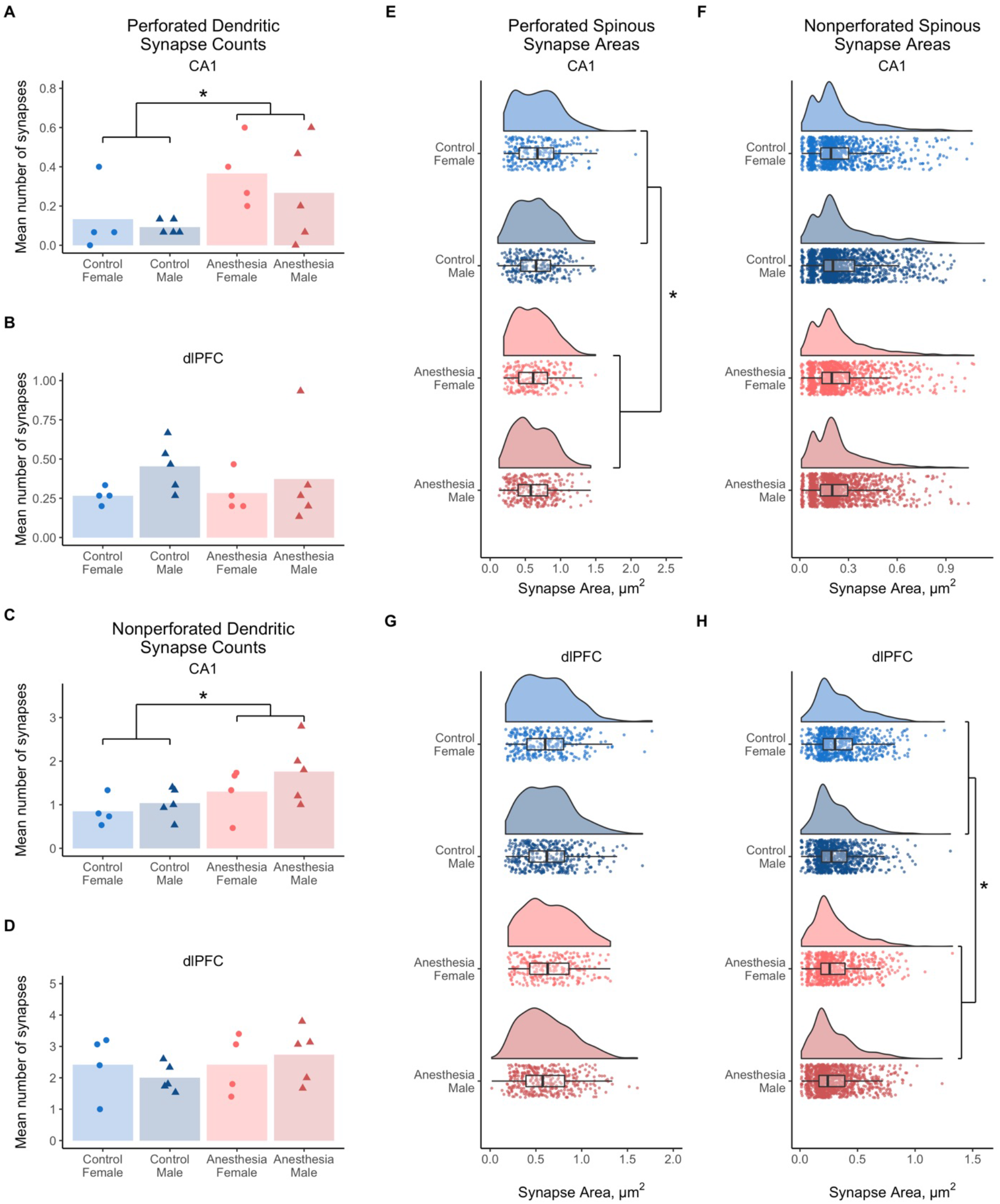
Synapse counts and areas vary by synapse type and region. (*A*) Monkeys exposed to anesthesia as infants had 180% more perforated dendritic synapses in CA1 than unexposed controls (χ2 (1) = 6.11, p = 0.013). (*B*) Numbers of perforated dendritic synapses in dlPFC are not affected by anesthesia (χ2 (1) = 0.22, p = 0.64), sex (χ2 (1) = 2.52, p = 0.11), or their interaction (χ2 (1) = 0.30, p = 0.58). (*C*) Monkeys exposed to anesthesia as infants had 63% more nonperforated dendritic synapses in CA1 than unexposed controls (χ2 (1) = 6.02, p = 0.014). (*D*) Numbers of nonperforated dendritic synapses in dlPFC are not affected by anesthesia (χ2 (1) = 1.13, p = 0.29), sex (χ2 (1) = 0.002, p = 0.96), or their interaction (χ2 (1) = 0.90, p = 0.34). (*E*) Perforated spinous synapse areas in CA1 were 7.5% lower in monkeys exposed to anesthesia in infancy compared to unexposed controls (F(1, 11.80) = 5.19, p = 0.042). (*F*) Areas of nonperforated spinous synapses in CA1 were not affected by anesthesia (F(1, 14.07) = 0.24, p = 0.63), sex (F(1, 14.07) = 0.05, p = 0.81), or their interaction (F(1, 14.07) = 0.94, p = 0.35). (*G*) Perforated spinous synapse areas in dlPFC trended toward a significant interaction of anesthesia and sex (F(1, 15.73) = 3.13, p = 0.096). (*H*) Areas of nonperforated spinous synapses in dlPFC are 10.4% lower in monkeys exposed to anesthesia in infancy compared to controls (F(1, 14.89) = 6.66, p = 0.021). (*A-H*) Unexposed controls shown in blue, monkeys exposed to anesthesia shown in red. In (*A-D*), females are lighter shades with circles, males are darker shades with triangles; individual points indicate individual monkey means. In (*E-H*), kernel distribution graphs show the relative spread of the frequency of areas, box & whisker plots show the range and quartiles of areas, and individual points are individual synapses. * = p < 0.05.

In the dlPFC, we also found significant interactions of treatment by sex (*χ*^2^(1) = 4.24, p = 0.04) and by synapse type (*χ*^2^(3) = 18.25, p = 0.0004) for numbers of synapses classified by target and complexity. However, no additional treatment, sex, or interaction effects on synapse numbers were found at the omnibus or individual category levels (p-values > 0.10), including for perforated and nonperforated dendritic synapses (**Figure 4B&D**). The contrast between the significant omnibus effects of early-life anesthesia on synapses by type and the absence of effects on individual categories of synapses could indicate that changes to the numbers of dlPFC synapses by anesthesia were more subtle than we could detect with our category sample sizes.

### Early-life anesthesia reduces perforated spinous synapse areas in CA1 and nonperforated spinous synapse areas in dlPFC

Our analyses of the areas of CA1 synapses categorized by target and complexity indicated significant interactions of anesthesia treatment with sex (F(1, 100.0) = 8.51, p = 0.004), including interactions with synapse target (F(1, 6720.5) = 13.13, p = 0.0003), synapse complexity (F(1, 6717.0) = 9.49, p = 0.002), and both target and complexity (F(1, 6717.3) = 5.31, p = 0.02). The combined interactions of anesthesia treatment with target, with complexity, and with both target and complexity significantly influenced synapse areas (*χ*^2^(3) = 8.71, p = 0.033). Likewise, synapse areas varied per the combined interactions of anesthetic treatment and sex together on synapse area by target, complexity, and target by complexity (*χ*^2^(3) = 18.97, p = 0.0003). The combined influence of sex interactions with target, complexity, and target & complexity on synapse area also trended toward significance (*χ*^2^(3) = 7.45, p = 0.059). These omnibus results showed that early-life anesthesia altered later-life synapse areas according to sex, postsynaptic target, and synapse complexity.

Our subsequent analyses of synapse subtypes revealed that the long-term effects of anesthesia on synapse areas in CA1 were limited to perforated spinous synapses. The mean area of perforated spinous synapses was 7.5% lower in monkeys exposed to anesthesia compared to controls (controls 0.67 ± 0.30 µm^2^, anesthesia 0.62 ± 0.27 µm^2^, F(1, 11.80) = 5.19, p = 0.042) (**Figure 4E**). No other treatment, sex, or interaction effects were found across individual synapse types, including nonperforated spinous synapses (p-values > 0.10) (**Figure 4F**). These results show that the long-term reduction in synaptic coverage in CA1 by early-life anesthesia was selective to perforated spinous synapses.

In the dlPFC, synapse areas were found to significantly vary by the sum of the interactions of anesthesia treatment and sex with synapse target, complexity, and target by complexity (*χ*^2^(3) = 8.77, p = 0.033). The sums of all such synapse type component interactions with treatment alone and with sex alone had no effects on synapse area in dlPFC (p-values > 0.20).

Individually, perforated spinous synapse areas in dlPFC showed a trend toward significance for the interaction of anesthesia and sex (F(1, 15.73) = 3.13, p = 0.096), with mean areas of 0.62 ± 0.28 and 0.64 ± 0.27 µm^2^ respectively for control females and males, and 0.65 ± 0.27 and 0.61 ± 0.28 µm^2^ respectively for females and males exposed to anesthesia as infants (**Figure 4G**).

Additionally, our analyses of individual synapse types showed that nonperforated spinous synapses in dlPFC were 10.4% smaller on average in monkeys exposed to anesthesia as infants compared to controls (F(1, 14.89) = 6.66, p = 0.021), and trended toward a significant effect of sex (F(1, 14.89) = 3.69, p = 0.074) (**Figure 4H**). Mean nonperforated spinous synapse areas were 0.35 ± 0.20 and 0.32 ± 0.18 µm^2^ respectively for control females and males, and 0.31 ± 0.19 and 0.29 ± 0.19 µm^2^ respectively for females and males exposed to anesthesia as infants. All other effects of anesthesia, sex, and their interaction did not affect dlPFC synapse areas by type (p-values > 0.10). These results indicated that anesthesia exposures in early life caused a specific long-term reduction to nonperforated spinous synapse areas in the dlPFC.

### CA1 presynaptic boutons show a specific reduction in curved mitochondria

When we measured the long-term impact of early-life anesthesia on the overall density of mitochondria within presynaptic boutons, we found that in CA1 and dlPFC alike, the numbers of mitochondria in presynaptic boutons per cubic micron of tissue were not affected by treatment, sex, or their interaction (p-values > 0.10) (**Figure 5A&C**). This showed that early-life exposures to anesthesia did not cause a gross, long-term change to presynaptic mitochondria in these two regions.

**Figure 5.**
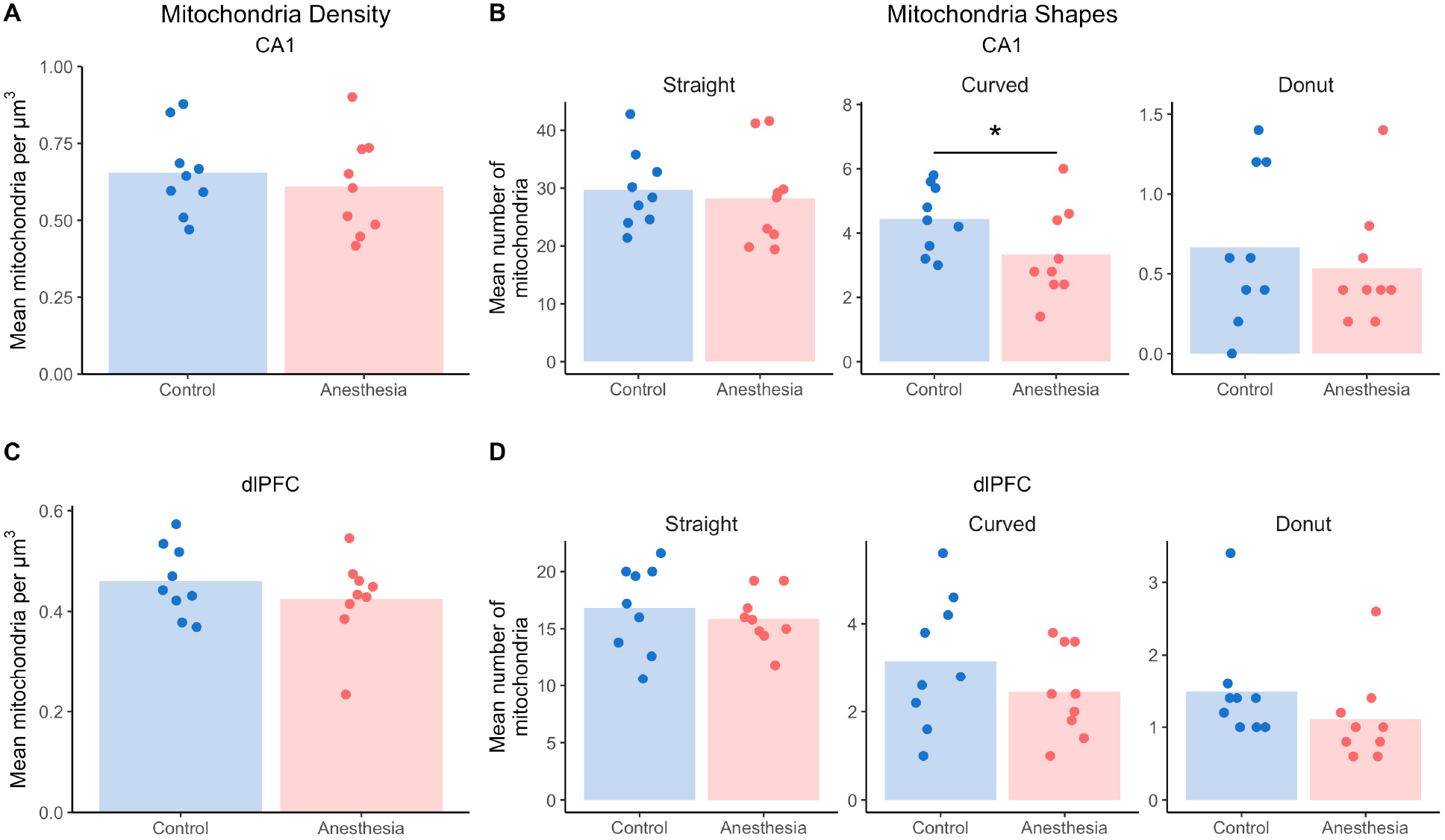
Early-life exposure to anesthesia selectively affected mitochondria shapes but not overall density mitochondria. (*A*) In CA1, the overall density of mitochondria in presynaptic boutons was not affected by exposure to early-life anesthesia (F(1, 14) = 0.72, p = 0.41), sex (F(1, 14) = 0.61, p = 0.45), of their interaction (F(1, 14) = 2.87, p = 0.11). (*B*) Monkeys exposed to anesthesia in infancy show a 25% reduction in curved mitochondria in CA1 compared to unexposed controls (χ2 (1= 4.12, p = 0.042), and no differences in straight or donut mitochondria (p values >0.15). (*C*) In dlPFC, the density of mitochondria in presynaptic boutons was not affected by early-life anesthesia (F(1, 14) = 1.10, p = 0.31), sex (F(1, 14) = 0.39, p = 0.54), or their interaction (F(1, 14) = 1.18, p = 0.30). (*D*) Exposure to anesthesia in early life did not alter numbers of straight (χ2 (1) = 0.33, p = 0.56), curved (χ2 (1) = 1.32, p = 0.25), or donut mitochondria (χ2 (1) = 1.59, p = 0.21). Unexposed controls shown in blue, monkeys exposed to anesthesia shown in red. Individual points indicate individual monkey means.

We also investigated whether early-life anesthesia had long-term effects on mitochondrial health and maintenance, as indicated by their straight, curved, or donut-shaped morphology. In CA1, monkeys that were exposed to anesthesia as infants had 25% fewer curved mitochondria in presynaptic boutons compared to controls (controls 4.44 ± 2.22, anesthesia 3.33 ± 2.14 mitochondria per volume, *χ*^2^(1) = 4.12, p = 0.042) (**Figure 5B**). There were no other differences in mitochondria morphology caused by anesthesia treatment, sex, or their interaction (p-values > 0.20). In the dlPFC, we found no effect of treatment, sex, or their interaction on mitochondria morphology (p-values > 0.10) (**Figure 5D**). In summary, exposures to anesthesia in infancy imparted a region-specific, long-term reduction in curved mitochondria in CA1 presynaptic boutons.

### Early-life anesthesia alters numbers of mitochondria contained by presynaptic boutons in dlPFC

Next, we assessed how early-life anesthesia affected the long-term, relative abundance of mitochondria within presynaptic boutons by measuring the numbers of presynaptic boutons containing 0, 1, 2, and 3 or more mitochondria. We found trends in CA1 toward the interaction of treatment and sex across all categories of mitochondria frequency within presynaptic boutons (*χ*^2^(1) = 2.87, p = 0.09), toward the interaction of treatment and sex for numbers of boutons with 0 mitochondria (*χ*^2^(1) = 3.19, p = 0.07), and toward a sex effect on boutons with 3 or more mitochondria (*χ*^2^(1) = 3.72, p = 0.05) (**Figure 6A**). No other treatment, sex, or interaction effects were observed at the global or individual levels in CA1 (p-values > 0.10). These results indicate that anesthesia exposures in early life did not significantly alter the long-term prevalence of mitochondria in presynaptic boutons in CA1.

**Figure 6.**
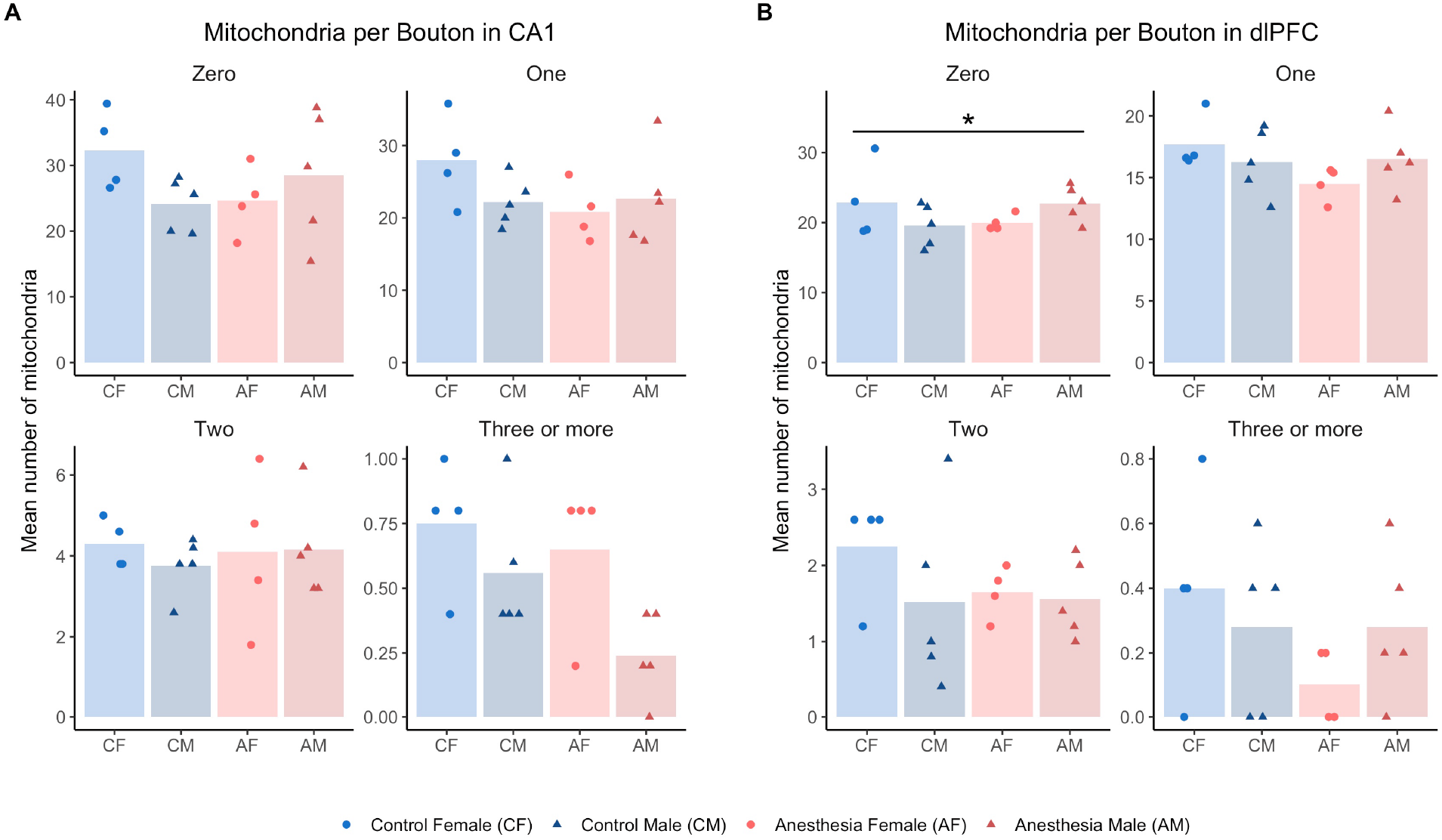
Early-life anesthesia exposure only significantly altered the numbers of mitochondria per presynaptic bouton in the dlPFC. (*A*) The numbers of mitochondria per presynaptic bouton in CA1 showed a trend toward a significant interaction of early-life anesthesia exposure with sex (χ2 (1) = 2.87, p = 0.09), and trends toward the interaction of anesthesia and sex for boutons with 0 mitochondria (*χ*^2^(1) = 3.19, p = 0.07), and sex for boutons with 3+ mitochondria (*χ*^2^(1) = 3.72, p = 0.05). (*B*) The numbers of mitochondria per presynaptic bouton in CA1 showed a significant interaction of early-life anesthesia exposure with sex (χ2 (1) = 6.13, p = 0.013), with an interaction of early-life anesthesia with sex for boutons with 0 mitochondria (χ2 (1) = 4.15, p = 0.042), and a trend toward an interaction effect of anesthesia and sex for boutons with 1 mitochondrion (χ2 (1) = 2.91, p = 0.088). Unexposed controls shown in blue, monkeys exposed to anesthesia shown in red. Females are lighter shades with circles, males are darker shades with triangles. Individual points indicate individual monkey means. * = p < 0.05 for individual categories.

In the dlPFC, the numbers of mitochondria per presynaptic bouton showed a significant treatment by sex interaction across categories (*χ*^2^(1) = 6.13, p = 0.013). There was a significant treatment by sex interaction in boutons with 0 mitochondria (*χ*^2^(1) = 4.15, p = 0.042) (**Figure 6B**), although posthoc pairwise comparisons were not significant (p-values > 0.05). Qualitatively, boutons with 0 mitochondria were decreased in females by anesthesia— 20 ± 3.9 boutons for females exposed to anesthesia compared to 22.8 ± 7.5 boutons for control females— and increased in males by anesthesia: 22.8 ± 5.5 boutons in males exposed to anesthesia compared to 19.6 ± 5.8 boutons for control males. Likewise, the numbers of boutons with 1 mitochondrion trended toward the opposite interaction effect of treatment and sex (*χ*^2^(1) = 2.91, p = 0.088), with more boutons in females exposed to anesthesia and fewer boutons in males exposed to anesthesia compared respectively to female and male controls. All other treatment, sex, and interaction effects in dlPFC were not significant (p-values > 0.10). These results demonstrate that the long-term impact of neonatal exposures to anesthesia on the prevalence of mitochondria in dlPFC presynaptic boutons is limited to boutons with a low frequency of mitochondria.

## DISCUSSION

We found that monkeys exposed to anesthesia multiple times as infants showed region-specific and sometimes sex-specific alterations in their synaptic ultrastructure 4 years later compared to unexposed monkeys. Monkeys exposed to anesthesia as infants showed a modulation of the areas of their largest synapses in CA1 and dlPFC, with sex-specific effects in CA1. When examining how individual subtypes of synapses contributed to these changes, we found that the area of perforated spinous synapses in CA1 was reduced by 7.6%, the area of nonperforated spinous synapses in dlPFC was reduced by 10.4%, and the numbers of dendritic shaft synapses in CA1 were elevated by 180% for perforated synapses and by 63% for nonperforated synapses in monkeys exposed to anesthesia in infancy. Likewise, monkeys exposed to anesthesia had a 25% reduction in curved mitochondria in CA1 presynaptic boutons, and a sex-specific modulation of the number of presynaptic boutons with 0 mitochondria in the dlPFC: a decrease for females and an increase for males.

### Long-term effects of anesthesia exposure on synapses

The loss of spinous synapse coverage in monkeys exposed to sevoflurane in infancy is consistent with effects of early-life anesthesia on spines in rodents. Similarly to our cohort, spinous synapse lengths were reduced in the CA1, 3-5 months after multiple neonatal sevoflurane exposures in male rats (Amrock et al., 2015). Spine head volumes correlate highly with synapse area (Harris & Weinberg, 2012), and also experience a downshift in their average size in rodent hippocampus and cortex after early-life anesthesia (Briner et al., 2010, 2011; De Roo et al., 2009). This downshift is characterized by both an outgrowth of smaller, labile spines and filopodia (Zimering et al., 2016) and the loss of large, stable mushroom spines (Kang et al., 2017; Schaefer et al., 2020; Zhou et al., 2019). Interestingly, the reduction in synapse areas that we found in 5-10% of the largest synapses in CA1 and dlPFC in absence of an overall change to synapse density suggests that a similar shift in spine morphology could be in effect years after anesthesia in nonhuman primates.

Likewise, the reduction in spinous synapse areas in CA1 and dlPFC of monkeys exposed to neonatal anesthesia implies an enduring loss of excitatory synapse coverage with specific long-term consequences for the makeup of excitatory receptors, and by extension excitatory neurotransmission, in both regions. Almost all (91-99%) spinous synapses in the CA1 and cortex of rodents and primates are excitatory, as demonstrated by their asymmetry (Blazquez-Llorca et al., 2021; Cano-Astorga et al., 2021; Domínguez-Álvaro et al., 2019; Hsu et al., 2017; Montero-Crespo et al., 2020; Santuy et al., 2018; Tao et al., 2018). Thus, the reductions in spinous synapse areas by early-life anesthesia in the present cohort of monkeys are likely highly specific to excitatory synapses. Moreover, in the adult rat CA1, 100% of perforated and 64% of nonperforated synapses contain AMPA receptors, and 100% of both subtypes contain NMDA receptors (Ganeshina et al., 2004a, 2004b). This suggests that the reduction in perforated synapse areas in CA1 and the reduction in nonperforated synapse areas in dlPFC could translate to AMPA and NMDA receptor loss in both regions. If the decrease in spinous synapse coverage in CA1 and dlPFC of monkeys does correspond to a diminution of AMPA and NMDA receptor numbers by anesthesia, then the observed decrease in spinous synapse areas could be a plausible substrate for the reduced excitatory postsynaptic potentials (Chung et al., 2017; Ju et al., 2019) and suppressed LTP shown after neonatal anesthesia exposure (Guo et al., 2018; Jevtovic-Todorovic et al., 2003; Kato et al., 2013; Schaefer et al., 2020; Sun et al., 2020).

On the other hand, the increase in perforated and nonperforated synapses targeting CA1 dendrites has more ambiguous implications. Dendritic synapses have been largely unexplored in the context of early-life anesthesia exposure, although one study in rats contrasted our finding, showing instead that a neonatal exposure to an anesthetic cocktail caused no difference in the ratio of mean spinous to dendritic synapse numbers in the subiculum two weeks later (Lunardi et al., 2010). However, subtle changes to dendritic synapses could be harder to detect in rats without targeted sampling because at baseline, synapses targeting dendrites comprise only 5% of the total synapses in rat CA1 (Blazquez-Llorca et al., 2021) compared to 21% in primate (human) CA1 (Montero-Crespo et al., 2020). In addition, dendritic synapses have a higher proportion of inhibitory symmetric synapses compared to spinous synapses—46% compared to <1%—in human stratum radiatum (Montero-Crespo et al., 2020). This suggests that the increase in dendritic synapses in monkeys by early-life anesthesia could demonstrate modulation of inhibitory or excitatory tone. Deeper exploration of the effects of early-life anesthesia on dendritic synapse populations will be needed to interrogate this result more fully.

### Long-term effects of anesthesia exposure on mitochondria

The alterations to presynaptic mitochondria in monkeys years after their anesthesia exposures in infancy extend our understanding of the impact of neonatal anesthesia on mitochondria from the rodent literature. The lack of long-term effects of anesthesia on the overall mitochondria density in CA1 and dlPFC presynaptic boutons in monkeys contrasts the acute and long-term modulation of mitochondria density by neonatal anesthesia in the subicular cytoplasm (Boscolo et al., 2013; Sanchez et al., 2011) and CA1 presynaptic terminals of rats (Amrock et al., 2015). The distinction between these findings could be due to any number of methodological differences that are common features of early-life anesthesia research, including and not limited to interspecies differences in neurodevelopment timelines, dose and type of anesthetics used, sex, and regions analyzed. Regardless, the lack of effect of anesthesia exposure on the overall mitochondria density in monkey CA1 and dlPFC suggests that mitochondria homeostasis dynamics are not grossly and enduringly impaired in these regions.

Rather, the reduction in curved mitochondria in the CA1 suggests the long-term effects of anesthesia on energy homeostasis in monkeys may take a more nuanced hue. The 25% decrease in curved mitochondria in CA1 may point to a disturbance in energy management processes tendered by mitochondrial fusion and fission. Curved mitochondria are frequently complex in shape and can form through the fusion of healthy mitochondria with damaged mitochondria to compensate for their dysfunctional components (Norat et al., 2020). This process unfolds over the course of postnatal development in rats (Smith et al., 2016), indicating mitochondrial fusion is a developmentally regulated process with a potential for susceptibility to insult during early life. Consistent with the acute disruption of the fusion-fission axis by anesthesia in rat pups (Boscolo et al., 2013) and neonatal rat hippocampal neurons and astrocytes (Y. Liu et al., 2017), the reduction in curved mitochondria we observed could be a function of decreased fusion, excessive fission, or both. Ultimately, the deficit in curved mitochondria could imply an ongoing complication in energy allocation for adolescent monkeys exposed to sevoflurane as infants that would likely be exacerbated during times of high energy demands, and could relate to their behavioral deficits.

The modulation of numbers of boutons with 0 mitochondria in dlPFC by early-life anesthesia hint at additional long-term changes in neuronal responsivity to energy needs. Healthy neurons at rest use 4.7 billion ATP per second (Zhu et al., 2012), and meet additional energy expenditure demands by recruiting mitochondria along filamentous actin (Li et al., 2020). However, it has been shown that anesthesia exposure can cause filamentous actin depolymerization (Platholi et al., 2014; Zimering et al., 2016), which could affect mitochondrial transport to presynaptic terminals. In addition, a decrease in mitochondrial membrane potential, such as that caused by anesthesia exposure (Bains et al., 2009), can trigger the phosphorylation and ubiquitination of the RhoGTPase Miro, which detaches mitochondria from the cytoskeleton and prevents their transport (Palikaras et al., 2018). Presynaptic boutons without mitochondria are reliant on ATP diffusion and show less vesicle docking in response to theta-burst stimulation that produces LTP (Smith et al., 2016). This suggests that the fluctuation in the number of presynaptic boutons lacking mitochondria in the dlPFC in monkeys exposed to anesthesia as infants could translate to variance in the synaptic efficiency of dlPFC neurons.

### Connecting structural changes to behavioral deficits

Notably, the impairments in synaptic plasticity and efficiency that are suggested by the synapse and mitochondria remodeling observed in monkeys exposed to anesthesia as infants are relevant to the memory deficits and socioemotional dysregulation seen in the same monkeys (Alvarado et al., 2017; Raper et al., 2015, 2018). In several mouse models of neurodevelopmental disorders, deficits in memory, social, and emotional behaviors have been linked to impairments in LTP that are related to synaptic restructuring in hippocampus and cortex (Belichenko et al., 2004; Katrancha et al., 2019; Uppal et al., 2015). Similarly, in mice exposed to ketamine as neonates, stressor-evoked anxiety-like behaviors in adulthood were linked to a deficit in LTP mediated by impaired AMPA recruitment to synapses (X. Zhang et al., 2020). The relationship between mitochondrial disturbances and behavioral perturbation has been further demonstrated by the correlations of synaptic mitochondrial health with working memory in monkeys and recognition memory in rats (Hara et al., 2014; Olesen et al., 2020), and the role of mitochondrial deficiency in generating anxiety-like behaviors in a chronic social defeat paradigm in mice (Duan et al., 2021). In all, these studies support a role for synaptic and mitochondrial remodeling in the behavioral deficits we previously observed in monkeys exposed to anesthesia as infants.

### Limitations

Although our sample size was relatively large for ultrastructural research in rhesus macaques, it was underpowered for evaluating sex-based effects, especially for modest effect sizes. We may have failed to detect more subtle treatment-by-sex effects, as suggested by trends toward significance in the numbers of dlPFC synapses by type and numbers of mitochondria in presynaptic boutons. These trends could relate to the evidence of sex, age, and anesthetic interaction effects on inhibitory and excitatory signaling that has emerged in rodents (Cabrera et al., 2020; Ju et al., 2019). Future experiments across ages and animal models will be instrumental in determining the extent that sex modulates the effects of neonatal anesthesia throughout the brain’s development.

In order to avoid confounding anesthesia effects, we avoided any experimental measures between anesthesia exposures and brain collection that would have required additional exposure to anesthesia, such as neuroimaging, electrophysiology, or additional tissue collection. As a result, we lack of structural or molecular data from the intervening years. Nonetheless, the ultrastructural changes we found four years after neonatal exposures to anesthesia provide novel insights into the long-lasting impacts of neonatal anesthesia on primate neurobiology.

Additionally, our anesthesia paradigm of three instances of four hours of sevoflurane may pertain best to children at the high end of anesthesia exposure. Approximately 26% of the 500,000 to 1 million infants and children exposed to anesthesia every year have a single exposure greater than 3 hours, or multiple exposures before age 3 (Ing et al., 2021; Shi et al., 2018). This means that the results we report here may be most relevant to those tens of thousands of infants annually with the greatest cumulative anesthetic history. Yet, because single exposures to anesthesia have also been found to confer an increased risk for behavioral problems in children (Ing et al., 2021), the structural findings we report here may also lend themselves to the long-term outcomes of shorter exposures to anesthesia.

## CONCLUSION

To our knowledge, this is the first study in nonhuman primates to show that exposure to general anesthesia in infancy leaves a structural signature on the brain that endures for years. Our findings extend prior research on the altering effects of early-life anesthesia on neuronal ultrastructure in rodents and nonhuman primates, and demonstrate that critical components of synapses in the hippocampus and prefrontal cortex of monkeys exposed to anesthesia in infancy retained deficits into adolescence. We suspect that the ultrastructural changes that we found in monkeys with neonatal anesthesia exposure may be related to their impairments in visual recognition memory and heightened emotional reactivity to social stressors.

In light of these findings, there are several lines of inquiry that could be explored in follow-up studies. To determine whether the reduction in synapse areas in CA1 and dlPFC signifies altered levels of excitatory receptors, we could perform immunogold staining for AMPA and NMDA receptors in tandem with ultrastructural analyses of synapses by type. In addition, we could apply our ultrastructural analyses to synapses and mitochondria in the pyramidal cell body layer of each region to further investigate the effects of early-life anesthesia on inhibitory synapses and mitochondria motility. Then, we could explore the long-term effects of neonatal anesthesia in other brain regions implicated in emotional regulation and memory. The cognitive control network has been implicated in aspects of emotional reactivity including attention shifting related to emotional and nonemotional stimuli, determining the valence of a social stimulus, and self-regulation (Perlman et al., 2014). Brain regions in this network, including the amygdala, anterior cingulate cortex, and orbitofrontal cortex, are thus an intriguing target for future queries of the link between the behavioral and ultrastructural outcomes of early-life anesthesia in later life.

## Acknowledgements

We thank Dr. Deanna Benson and Dr. George Huntley for their helpful suggestions on experimental design. We thank Allison Sowa for her help with electron microscopy protocols, and Camille Casiño for reviewing the manuscript. This research was supported by a 2016 NSF Graduate Research Fellowship (TF), and NICHD grants R01-HD068388 and R01-HD099231 (MGB).

---

**Supplemental table:**
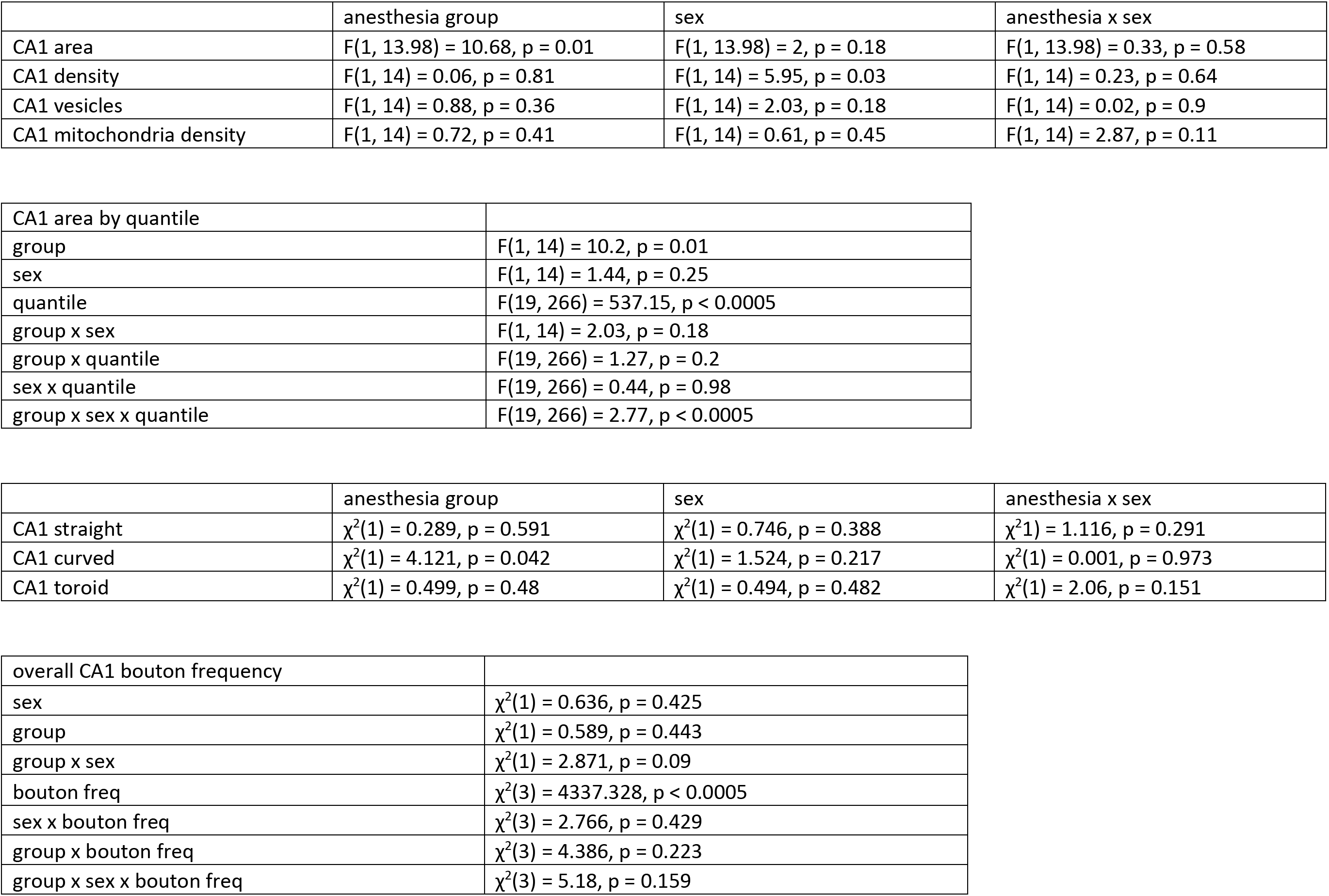
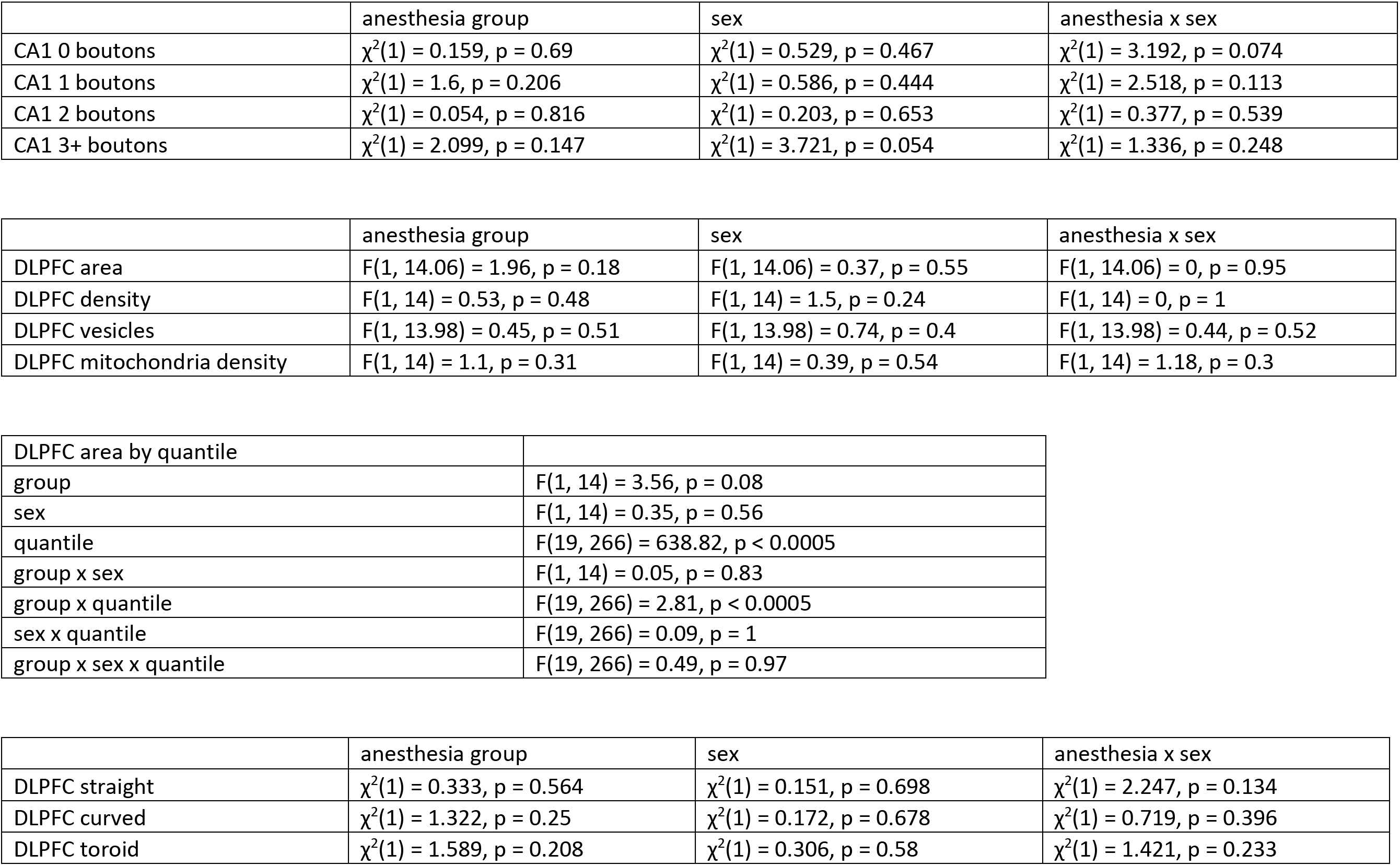
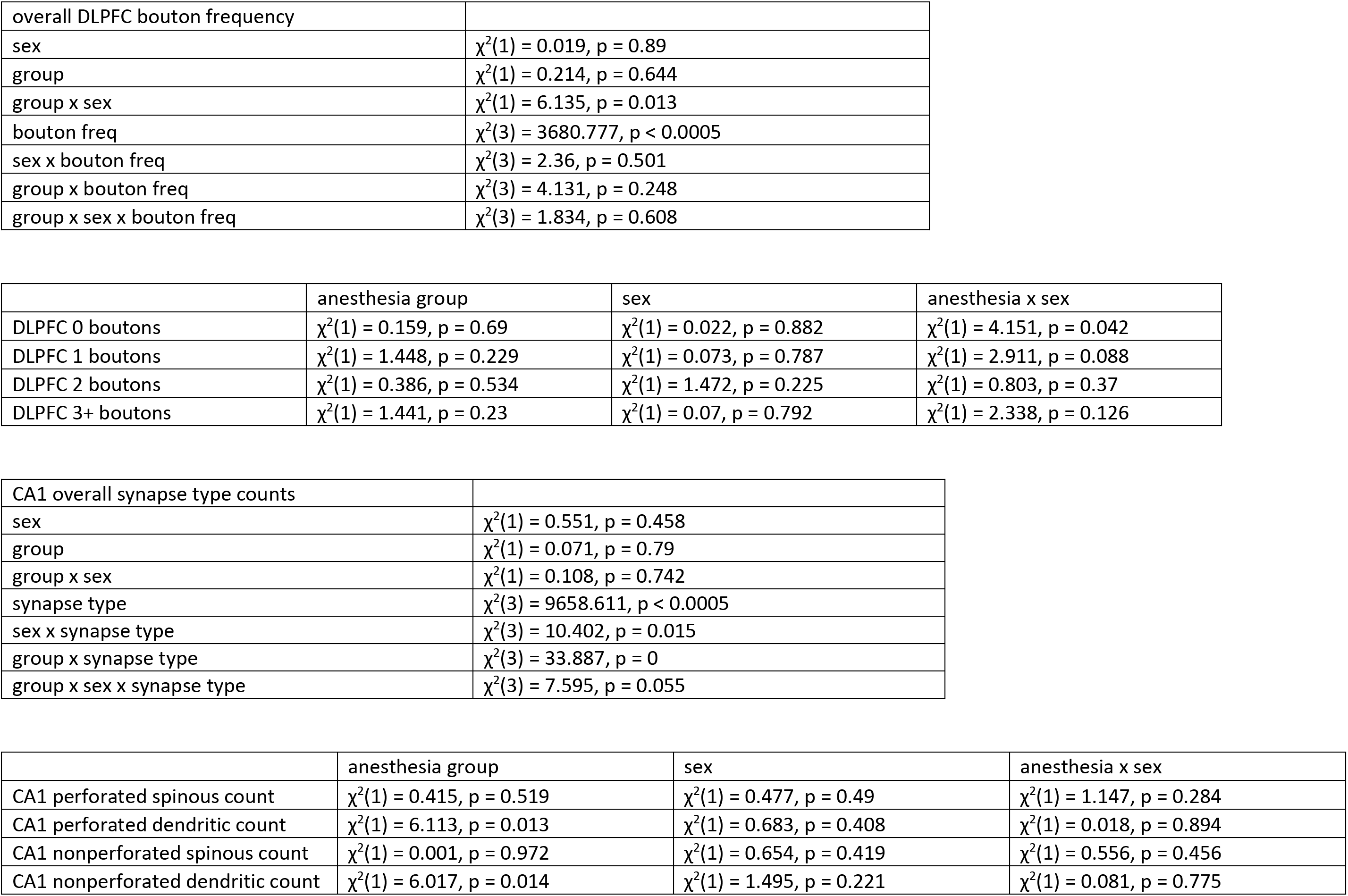
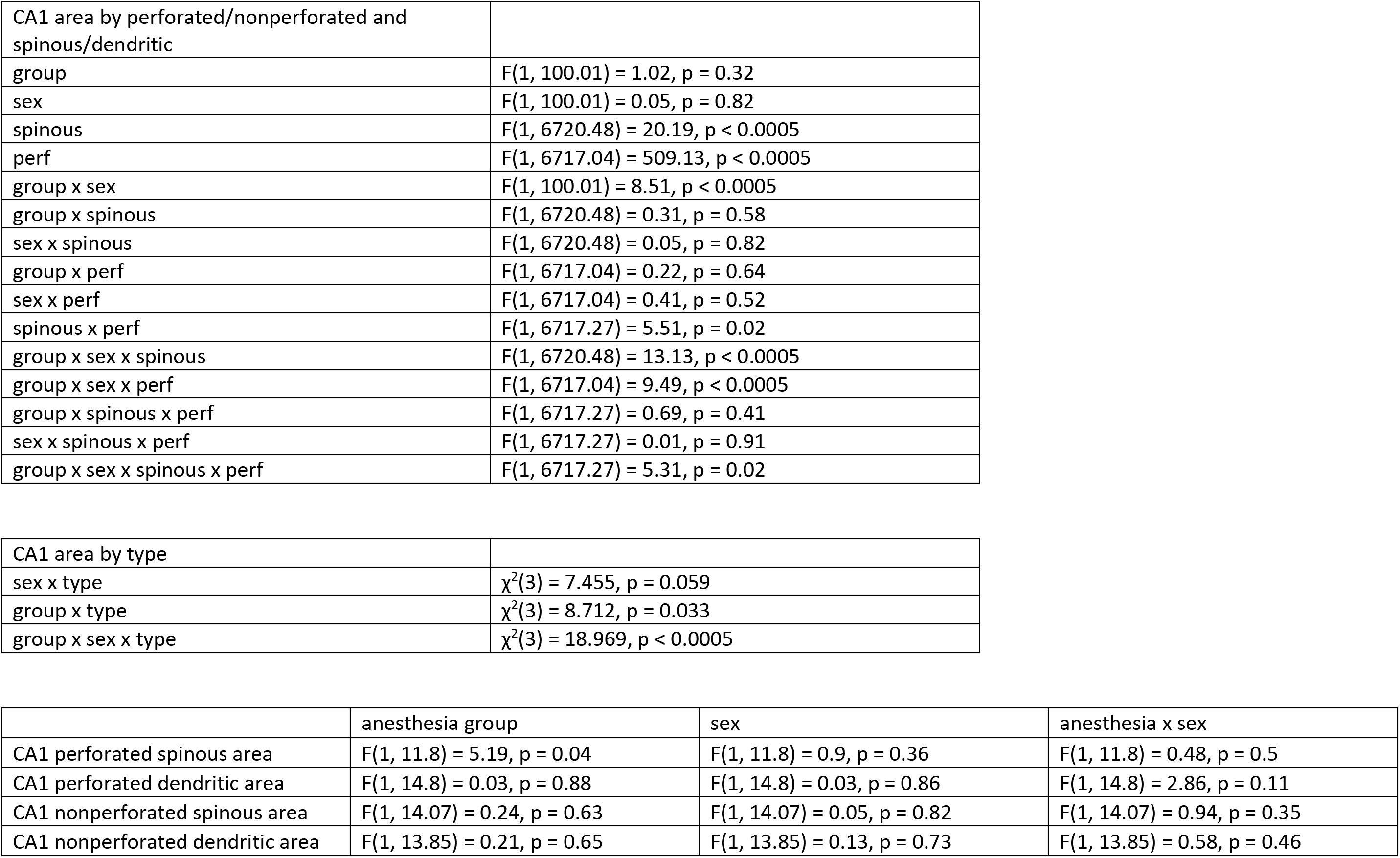
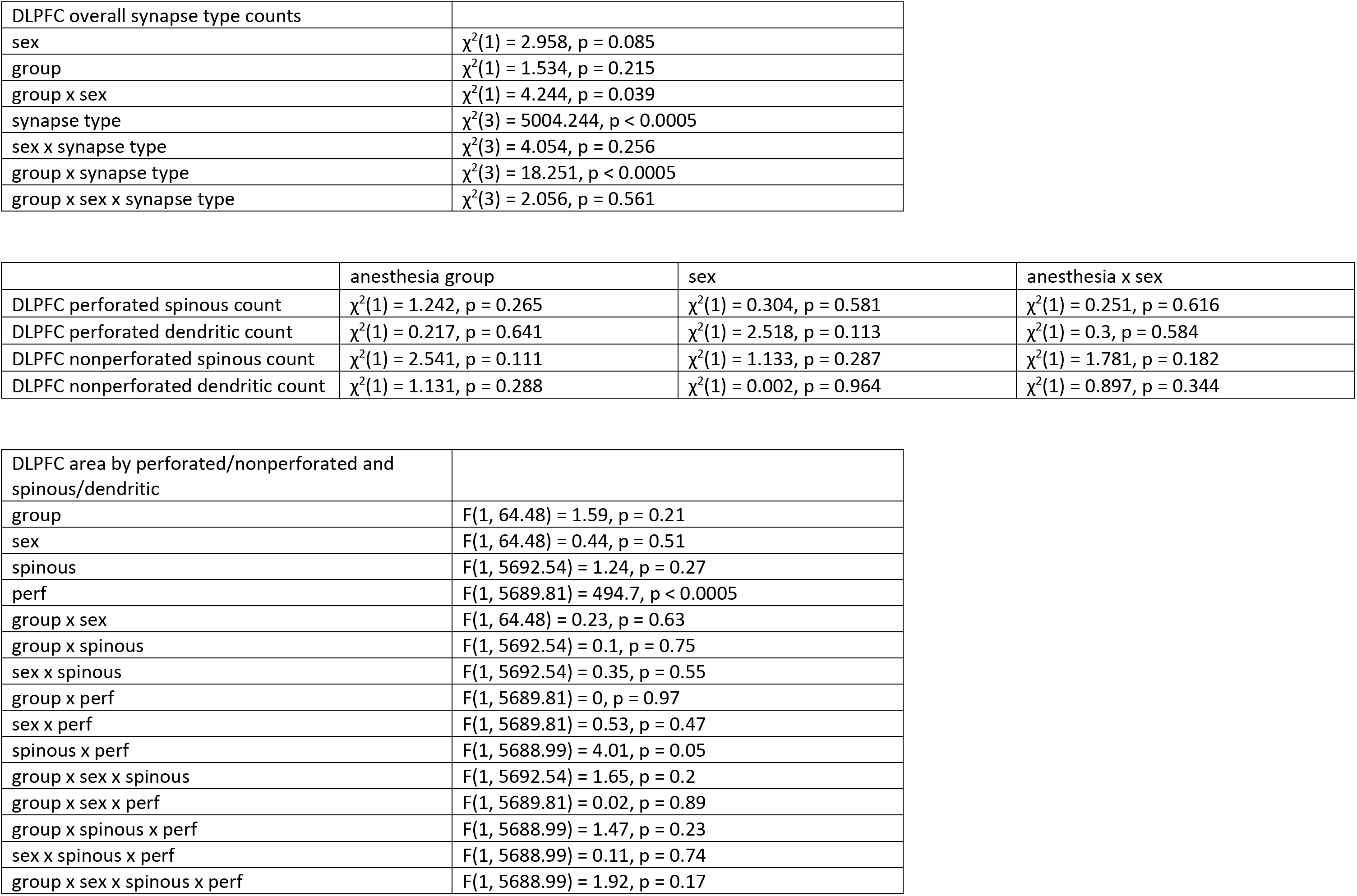
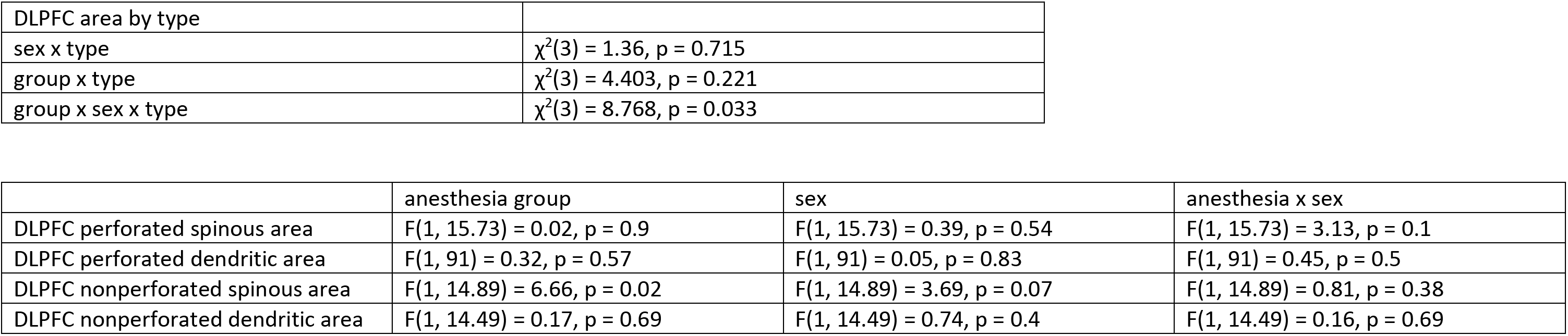
Statistical summaries of all analyses.

|  | anesthesia group | sex | anesthesia x sex |
| --- | --- | --- | --- |
| CA1 area | $F(1, 13.98) = 10.68, p = 0.01$ | $F(1, 13.98) = 2, p = 0.18$ | $F(1, 13.98) = 0.33, p = 0.58$ |
| CA1 density | $F(1, 14) = 0.06, p = 0.81$ | $F(1, 14) = 5.95, p = 0.03$ | $F(1, 14) = 0.23, p = 0.64$ |
| CA1 vesicles | $F(1, 14) = 0.88, p = 0.36$ | $F(1, 14) = 2.03, p = 0.18$ | $F(1, 14) = 0.02, p = 0.9$ |
| CA1 mitochondria density | $F(1, 14) = 0.72, p = 0.41$ | $F(1, 14) = 0.61, p = 0.45$ | $F(1, 14) = 2.87, p = 0.11$ |

|  |  |
| --- | --- |
| CA1 area by quantile |  |
| group | $F(1, 14) = 10.2, p = 0.01$ |
| sex | $F(1, 14) = 1.44, p = 0.25$ |
| quantile | $F(19, 266) = 537.15, p < 0.0005$ |
| group x sex | $F(1, 14) = 2.03, p = 0.18$ |
| group x quantile | $F(19, 266) = 1.27, p = 0.2$ |
| sex x quantile | $F(19, 266) = 0.44, p = 0.98$ |
| group x sex x quantile | $F(19, 266) = 2.77, p < 0.0005$ |

|  | anesthesia group | sex | anesthesia x sex |
| --- | --- | --- | --- |
| CA1 straight | $\chi^2(1) = 0.289, p = 0.591$ | $\chi^2(1) = 0.746, p = 0.388$ | $\chi^2(1) = 1.116, p = 0.291$ |
| CA1 curved | $\chi^2(1) = 4.121, p = 0.042$ | $\chi^2(1) = 1.524, p = 0.217$ | $\chi^2(1) = 0.001, p = 0.973$ |
| CA1 toroid | $\chi^2(1) = 0.499, p = 0.48$ | $\chi^2(1) = 0.494, p = 0.482$ | $\chi^2(1) = 2.06, p = 0.151$ |

|  |  |
| --- | --- |
| overall CA1 bouton frequency |  |
| sex | $\chi^2(1) = 0.636, p = 0.425$ |
| group | $\chi^2(1) = 0.589, p = 0.443$ |
| group x sex | $\chi^2(1) = 2.871, p = 0.09$ |
| bouton freq | $\chi^2(3) = 4337.328, p < 0.0005$ |
| sex x bouton freq | $\chi^2(3) = 2.766, p = 0.429$ |
| group x bouton freq | $\chi^2(3) = 4.386, p = 0.223$ |
| group x sex x bouton freq | $\chi^2(3) = 5.18, p = 0.159$ |
| CA1 0 boutons | $\chi^2(1) = 0.159, p = 0.69$ | $\chi^2(1) = 0.529, p = 0.467$ | $\chi^2(1) = 3.192, p = 0.074$ |
| CA1 1 boutons | $\chi^2(1) = 1.6, p = 0.206$ | $\chi^2(1) = 0.586, p = 0.444$ | $\chi^2(1) = 2.518, p = 0.113$ |
| CA1 2 boutons | $\chi^2(1) = 0.054, p = 0.816$ | $\chi^2(1) = 0.203, p = 0.653$ | $\chi^2(1) = 0.377, p = 0.539$ |
| CA1 3+ boutons | $\chi^2(1) = 2.099, p = 0.147$ | $\chi^2(1) = 3.721, p = 0.054$ | $\chi^2(1) = 1.336, p = 0.248$ |
| DLPFC area | $F(1, 14.06) = 1.96, p = 0.18$ | $F(1, 14.06) = 0.37, p = 0.55$ | $F(1, 14.06) = 0, p = 0.95$ |
| DLPFC density | $F(1, 14) = 0.53, p = 0.48$ | $F(1, 14) = 1.5, p = 0.24$ | $F(1, 14) = 0, p = 1$ |
| DLPFC vesicles | $F(1, 13.98) = 0.45, p = 0.51$ | $F(1, 13.98) = 0.74, p = 0.4$ | $F(1, 13.98) = 0.44, p = 0.52$ |
| DLPFC mitochondria density | $F(1, 14) = 1.1, p = 0.31$ | $F(1, 14) = 0.39, p = 0.54$ | $F(1, 14) = 1.18, p = 0.3$ |

| DLPFC area by quantile |  |
| --- | --- |
| group | $F(1, 14) = 3.56, p = 0.08$ |
| sex | $F(1, 14) = 0.35, p = 0.56$ |
| quantile | $F(19, 266) = 638.82, p < 0.0005$ |
| group x sex | $F(1, 14) = 0.05, p = 0.83$ |
| group x quantile | $F(19, 266) = 2.81, p < 0.0005$ |
| sex x quantile | $F(19, 266) = 0.09, p = 1$ |
| group x sex x quantile | $F(19, 266) = 0.49, p = 0.97$ |

|  | anesthesia group | sex | anesthesia x sex |
| --- | --- | --- | --- |
| DLPFC straight | $\chi^2(1) = 0.333, p = 0.564$ | $\chi^2(1) = 0.151, p = 0.698$ | $\chi^2(1) = 2.247, p = 0.134$ |
| DLPFC curved | $\chi^2(1) = 1.322, p = 0.25$ | $\chi^2(1) = 0.172, p = 0.678$ | $\chi^2(1) = 0.719, p = 0.396$ |
| DLPFC toroid | $\chi^2(1) = 1.589, p = 0.208$ | $\chi^2(1) = 0.306, p = 0.58$ | $\chi^2(1) = 1.421, p = 0.233$ |

|  |  |
| --- | --- |
| overall DLPFC bouton frequency |  |
| sex | $\chi^2(1) = 0.019, p = 0.89$ |
| group | $\chi^2(1) = 0.214, p = 0.644$ |
| group x sex | $\chi^2(1) = 6.135, p = 0.013$ |
| bouton freq | $\chi^2(3) = 3680.777, p < 0.0005$ |
| sex x bouton freq | $\chi^2(3) = 2.36, p = 0.501$ |
| group x bouton freq | $\chi^2(3) = 4.131, p = 0.248$ |
| group x sex x bouton freq | $\chi^2(3) = 1.834, p = 0.608$ |

|  | anesthesia group | sex | anesthesia x sex |
| --- | --- | --- | --- |
| DLPFC 0 boutons | $\chi^2(1) = 0.159, p = 0.69$ | $\chi^2(1) = 0.022, p = 0.882$ | $\chi^2(1) = 4.151, p = 0.042$ |
| DLPFC 1 boutons | $\chi^2(1) = 1.448, p = 0.229$ | $\chi^2(1) = 0.073, p = 0.787$ | $\chi^2(1) = 2.911, p = 0.088$ |
| DLPFC 2 boutons | $\chi^2(1) = 0.386, p = 0.534$ | $\chi^2(1) = 1.472, p = 0.225$ | $\chi^2(1) = 0.803, p = 0.37$ |
| DLPFC 3+ boutons | $\chi^2(1) = 1.441, p = 0.23$ | $\chi^2(1) = 0.07, p = 0.792$ | $\chi^2(1) = 2.338, p = 0.126$ |

|  |  |
| --- | --- |
| CA1 overall synapse type counts |  |
| sex | $\chi^2(1) = 0.551, p = 0.458$ |
| group | $\chi^2(1) = 0.071, p = 0.79$ |
| group x sex | $\chi^2(1) = 0.108, p = 0.742$ |
| synapse type | $\chi^2(3) = 9658.611, p < 0.0005$ |
| sex x synapse type | $\chi^2(3) = 10.402, p = 0.015$ |
| group x synapse type | $\chi^2(3) = 33.887, p = 0$ |
| group x sex x synapse type | $\chi^2(3) = 7.595, p = 0.055$ |

|  | anesthesia group | sex | anesthesia x sex |
| --- | --- | --- | --- |
| CA1 perforated spinous count | $\chi^2(1) = 0.415, p = 0.519$ | $\chi^2(1) = 0.477, p = 0.49$ | $\chi^2(1) = 1.147, p = 0.284$ |
| CA1 perforated dendritic count | $\chi^2(1) = 6.113, p = 0.013$ | $\chi^2(1) = 0.683, p = 0.408$ | $\chi^2(1) = 0.018, p = 0.894$ |
| CA1 nonperforated spinous count | $\chi^2(1) = 0.001, p = 0.972$ | $\chi^2(1) = 0.654, p = 0.419$ | $\chi^2(1) = 0.556, p = 0.456$ |
| CA1 nonperforated dendritic count | $\chi^2(1) = 6.017, p = 0.014$ | $\chi^2(1) = 1.495, p = 0.221$ | $\chi^2(1) = 0.081, p = 0.775$ |

|  |  |
| --- | --- |
| CA1 area by perforated/nonperforated and spinous/dendritic |  |
| group | $F(1, 100.01) = 1.02, p = 0.32$ |
| sex | $F(1, 100.01) = 0.05, p = 0.82$ |
| spinous | $F(1, 6720.48) = 20.19, p < 0.0005$ |
| perf | $F(1, 6717.04) = 509.13, p < 0.0005$ |
| group x sex | $F(1, 100.01) = 8.51, p < 0.0005$ |
| group x spinous | $F(1, 6720.48) = 0.31, p = 0.58$ |
| sex x spinous | $F(1, 6720.48) = 0.05, p = 0.82$ |
| group x perf | $F(1, 6717.04) = 0.22, p = 0.64$ |
| sex x perf | $F(1, 6717.04) = 0.41, p = 0.52$ |
| spinous x perf | $F(1, 6717.27) = 5.51, p = 0.02$ |
| group x sex x spinous | $F(1, 6720.48) = 13.13, p < 0.0005$ |
| group x sex x perf | $F(1, 6717.04) = 9.49, p < 0.0005$ |
| group x spinous x perf | $F(1, 6717.27) = 0.69, p = 0.41$ |
| sex x spinous x perf | $F(1, 6717.27) = 0.01, p = 0.91$ |
| group x sex x spinous x perf | $F(1, 6717.27) = 5.31, p = 0.02$ |
| CA1 area by type |  |
| sex x type | $\chi^2(3) = 7.455, p = 0.059$ |
| group x type | $\chi^2(3) = 8.712, p = 0.033$ |
| group x sex x type | $\chi^2(3) = 18.969, p < 0.0005$ |

|  | anesthesia group | sex | anesthesia x sex |
| --- | --- | --- | --- |
| CA1 perforated spinous area | $F(1, 11.8) = 5.19, p = 0.04$ | $F(1, 11.8) = 0.9, p = 0.36$ | $F(1, 11.8) = 0.48, p = 0.5$ |
| CA1 perforated dendritic area | $F(1, 14.8) = 0.03, p = 0.88$ | $F(1, 14.8) = 0.03, p = 0.86$ | $F(1, 14.8) = 2.86, p = 0.11$ |
| CA1 nonperforated spinous area | $F(1, 14.07) = 0.24, p = 0.63$ | $F(1, 14.07) = 0.05, p = 0.82$ | $F(1, 14.07) = 0.94, p = 0.35$ |
| CA1 nonperforated dendritic area | $F(1, 13.85) = 0.21, p = 0.65$ | $F(1, 13.85) = 0.13, p = 0.73$ | $F(1, 13.85) = 0.58, p = 0.46$ |

|  |  |
| --- | --- |
| DLPFC overall synapse type counts |  |
| sex | $\chi^2(1) = 2.958, p = 0.085$ |
| group | $\chi^2(1) = 1.534, p = 0.215$ |
| group x sex | $\chi^2(1) = 4.244, p = 0.039$ |
| synapse type | $\chi^2(3) = 5004.244, p < 0.0005$ |
| sex x synapse type | $\chi^2(3) = 4.054, p = 0.256$ |
| group x synapse type | $\chi^2(3) = 18.251, p < 0.0005$ |
| group x sex x synapse type | $\chi^2(3) = 2.056, p = 0.561$ |

|  | anesthesia group | sex | anesthesia x sex |
| --- | --- | --- | --- |
| DLPFC perforated spinous count | $\chi^2(1) = 1.242, p = 0.265$ | $\chi^2(1) = 0.304, p = 0.581$ | $\chi^2(1) = 0.251, p = 0.616$ |
| DLPFC perforated dendritic count | $\chi^2(1) = 0.217, p = 0.641$ | $\chi^2(1) = 2.518, p = 0.113$ | $\chi^2(1) = 0.3, p = 0.584$ |
| DLPFC nonperforated spinous count | $\chi^2(1) = 2.541, p = 0.111$ | $\chi^2(1) = 1.133, p = 0.287$ | $\chi^2(1) = 1.781, p = 0.182$ |
| DLPFC nonperforated dendritic count | $\chi^2(1) = 1.131, p = 0.288$ | $\chi^2(1) = 0.002, p = 0.964$ | $\chi^2(1) = 0.897, p = 0.344$ |

|  |  |
| --- | --- |
| DLPFC area by perforated/nonperforated and spinous/dendritic |  |
| group | $F(1, 64.48) = 1.59, p = 0.21$ |
| sex | $F(1, 64.48) = 0.44, p = 0.51$ |
| spinous | $F(1, 5692.54) = 1.24, p = 0.27$ |
| perf | $F(1, 5689.81) = 494.7, p < 0.0005$ |
| group x sex | $F(1, 64.48) = 0.23, p = 0.63$ |
| group x spinous | $F(1, 5692.54) = 0.1, p = 0.75$ |
| sex x spinous | $F(1, 5692.54) = 0.35, p = 0.55$ |
| group x perf | $F(1, 5689.81) = 0, p = 0.97$ |
| sex x perf | $F(1, 5689.81) = 0.53, p = 0.47$ |
| spinous x perf | $F(1, 5688.99) = 4.01, p = 0.05$ |
| group x sex x spinous | $F(1, 5692.54) = 1.65, p = 0.2$ |
| group x sex x perf | $F(1, 5689.81) = 0.02, p = 0.89$ |
| group x spinous x perf | $F(1, 5688.99) = 1.47, p = 0.23$ |
| sex x spinous x perf | $F(1, 5688.99) = 0.11, p = 0.74$ |
| group x sex x spinous x perf | $F(1, 5688.99) = 1.92, p = 0.17$ |
| DLPFC area by type |  |
| sex x type | $\chi^2(3) = 1.36, p = 0.715$ |
| group x type | $\chi^2(3) = 4.403, p = 0.221$ |
| group x sex x type | $\chi^2(3) = 8.768, p = 0.033$ |

|  | anesthesia group | sex | anesthesia x sex |
| --- | --- | --- | --- |
| DLPFC perforated spinous area | $F(1, 15.73) = 0.02, p = 0.9$ | $F(1, 15.73) = 0.39, p = 0.54$ | $F(1, 15.73) = 3.13, p = 0.1$ |
| DLPFC perforated dendritic area | $F(1, 91) = 0.32, p = 0.57$ | $F(1, 91) = 0.05, p = 0.83$ | $F(1, 91) = 0.45, p = 0.5$ |
| DLPFC nonperforated spinous area | $F(1, 14.89) = 6.66, p = 0.02$ | $F(1, 14.89) = 3.69, p = 0.07$ | $F(1, 14.89) = 0.81, p = 0.38$ |
| DLPFC nonperforated dendritic area | $F(1, 14.49) = 0.17, p = 0.69$ | $F(1, 14.49) = 0.74, p = 0.4$ | $F(1, 14.49) = 0.16, p = 0.69$ |

